# REAL-TIME VISUALIZATION OF SPLICEOSOME ASSEMBLY REVEALS BASIC PRINCIPLES OF SPLICE SITE SELECTION

**DOI:** 10.1101/2024.07.12.603320

**Authors:** Benjamin T Donovan, Bixuan Wang, Gloria R Garcia, Stephen M Mount, Daniel R Larson

## Abstract

The spliceosome is a megadalton protein-RNA complex which removes introns from pre-mRNA, yet the dynamic early assembly steps have not been structurally resolved. Specifically, how the spliceosome selects the correct 3’ splice site (3’SS) amongst highly similar non-functional sites is not known. Here, we develop a kinetic model of splice site selection based on single-molecule U2AF heterodimer imaging *in vitro* and *in vivo*. The model successfully predicts alternative splicing patterns and indicates that 3’SS selection occurs while U2AF is in complex with the spliceosome, not during initial binding. This finding indicates the spliceosome operates in a ‘partial’ kinetic proofreading regime, catalyzed in part by the helicase DDX42, which increases selectivity to the underlying U2AF binding site while still allowing for efficient forward progression.

**ONE-SENTENCE SUMMARY:** We apply a kinetic proofreading model to elucidate how transient U2AF binding leads to high fidelity splice site selection.

## MAIN TEXT

RNA splicing is catalyzed by the spliceosome, a megadalton enzyme (*1*) consisting of RNA and hundreds of protein components, each serving a role in regulating splicing accuracy and speed (*2*). This complexity enables very fine regulation as nearly all human pre-mRNAs undergo some form of alternative splicing (*3*). Additionally, despite being a single turn-over enzyme, the spliceosome also exhibits exceptionally high enzymatic activity; recent work indicates that a single intron may be removed in multiple splicing reactions, through a process called recursive splicing (*4–7*). While both alternative and recursive splicing are regulated at the step of initial splice site recognition, there is no unifying mechanism to describe the basis of splice site choice in these different contexts. Moreover, despite the tremendous advances in our structural understanding of the spliceosome (*8–11*), many aspects of the early steps of spliceosome assembly remain unresolved, likely due to the transient nature of these complexes. Here, we use a combination of equilibrium binding assays and time-resolved kinetic assays both *in vitro* and *in vivo* to elucidate splice site selection and splicing fidelity.

### 3’ SPLICE SITES (3’SSs) CONTAIN HIGHLY DEGENERATE U2AF BINDING SITES

The early steps of splicing require recognition of the *cis*-acting sequences in pre-mRNA such as the 5’ splice site (**5’SS**), the branchpoint (**bp**), and the 3’ splice site (**3’SS**) by RNA binding proteins and snRNPs. Here, we focus on the kinetics of 3’SS selection by the U2AF complex. While the 5’SS is recognized directly by the U1 snRNP, the U2 snRNP is recruited to the 3’SS by RNA binding proteins including the essential U2AF heterodimer which is involved in nearly all splicing reactions. U2AF is comprised of U2AF2 and U2AF1 which bind the pyrimidine tract and the -AG dinucleotide that marks the 3’SS, respectively (*12–14*). U2AF plays a fundamental role in establishing the 3’SS because it is involved in spliceosome progression from initial splice site recognition in the early (E) complex to A complex assembly (*15*). Despite the paramount importance of accurate splicing, U2AF binds promiscuously throughout a pre-mRNA (*6*, *16*, *17*). Therefore, we set out to understand the characteristics of U2AF binding that promote splicing progression at a specific site but not others.

We addressed this question first with RNA Bind-N-Seq (RBNS), a high-throughput *in vitro* binding assay to quantitatively measure U2AF binding preferences to every RNA 12mer (*18*, *19*) (**Fig. S1A**). In contrast to previous SELEX assays for U2AF (*12*, *13*), we reasoned that RBNS would provide new insight into the U2AF binding landscape to all potential sites, including weak and moderate affinity sites which may be lost after multiple rounds of enrichment required for SELEX. In this assay, full-length U2AF heterodimer purified from insect cells (**Fig. 1A**) is mixed with a random pool of RNAs. After equilibration, RNAs in complex with U2AF are isolated by U2AF1-FLAG immunoprecipitation (IP) and deep sequenced (***see methods***). We find that the best predictor of enrichment for a specific RNA is the number of uridines in the binding site (**Fig. 1B**), indicating that U2AF2 is the main determinant of overall binding affinity. Additionally, we find that the contribution of U2AF1 to overall binding affinity is small: it is more advantageous to increase the length of the pyrimidine tract than to add an AG dinucleotide. As such, the majority of significantly enriched RNAs do not contain an AG (**Fig. 1C**). We do, however, detect an AG- preference when comparing an AG-containing sequence to an otherwise identical GA-containing sequence when the AG is at the 3’ end of the binding site (**Fig. 1D**).

**Figure 1:**
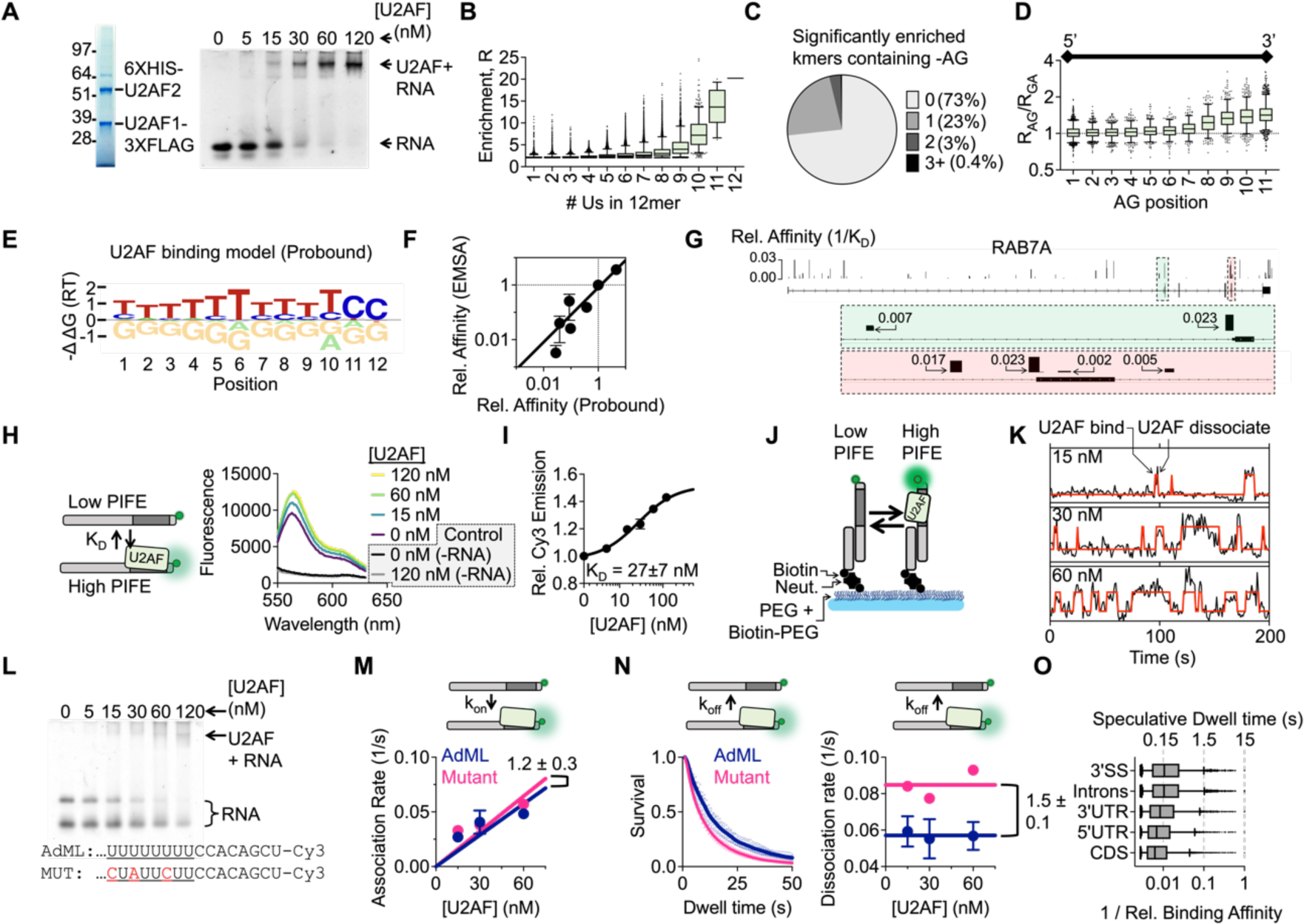
In vitro characterization of U2AF binding transcriptome wide A. Full length U2AF was purified from insect cells. We tested functionality by electrophoresis mobility shift assay (EMSA). B. Measuring U2AF binding to all possible 12mers in parallel with RNA Bind-N-Seq. U2AF heterodimer was mixed with a random pool of RNAs. Bound RNAs were isolated by U2AF1- FLAG IP and sequenced. The plot displays the distribution enrichment (R = Freq_IP_ /Freq_input_) segregated by number of uridines in 12mer (this data represents enrichment for 20 nM U2AF, n=2 replicates). C. The distribution of significantly enriched RNAs containing -AG dinucleotide. D. The enrichment ratio of an –AG containing sequence relative to an otherwise identical –GA containing sequence segregated by position of the -AG/-GA in the 12mer. E. Prediction of free energy contribution of each nucleotide using probound. F. Validating probound with competitive binding assays resolved by EMSA (see **Fig. S2**). Error bars represent one standard devation. G. Assigning U2AF binding affinity to *in vivo* binding sites detected with PAR-CLIP. In the browser shot of RAB7A, the height of each bar represents binding affinity relative to the sequence in (E). H. Adopting the PIFE assay to detect U2AF binding *in vitro* by bulk fluorescence. U2AF binding results in a ∼1.5 fold increase in fluorescence (n=2 replicates). As a control, we also performed the assay in the absence of RNA and find that intrinsic U2AF autofluorescence contributes very little to measured emission. I. Plotting relative change in fluorescence when titrating U2AF. Fitting with non-cooperative Hill Function (K_D_ = 27 ± 7 nM) agrees with EMSA measurements. Error bars represent one standard deviation. J. Applying the PIFE assay to the single-molecule level. RNAs are tethered to glass microscope slide through biotin-neutravidin linkage. Single surface-tethered RNAs are illuminated by Total Internal Fluorescence Reflection (TIRF) microscopy. K. U2AF binding and dissociation on individual RNAs is inferred from two-state Hidden Markov Model. L. EMSA of U2AF binding to an RNA with a mutated pyrimidine tract. M. U2AF binding rate to the AdML and mutant 3’ SS (k_on AdML_ = 0.0010 ± 0.0001 nM^-1^ s^-1^, k_on Mutant_ = 0.0011 ± 0.0002 nM^-1^ s^-1^) (n = 3 replicates for AdML, n=2 replicates for Mutant 3’SS). Error bars represent one standard deviation. N. Left: plotting U2AF dwell times for all replicates to AdML (n=9 measurements) and Mutant (n=6 measurements) 3’SS. Dotted lines represent individual replicates and solid lines represent average of all replicates. Right: Comparing U2AF dissociation rate from the AdML and mutant 3’SS at three concentrations of U2AF. U2AF dissociates about 1.5-fold faster from the mutant 3’SS (k_off AdML_ = 0.057 ± 0.003 s^-1^, k_off Mutant_ = 0.085 ± 0.003 s^-1^). Error bars represent one standard deviation. O. Comparing the distribution of U2AF binding affinities (relative to consensus sequence in Fig. 1E) for different binding site types. This plot only contains binding affinities within the range that were validated in Fig. 1F (>370-fold weaker than consensus). A speculative range of dwell times for these interactions is included on the top x-axis.

Next, we used the computational model Probound (*20*) to predict U2AF binding affinity to each site identified from a previous PAR-CLIP experiment (*6*) using the *in vitro* RBNS data (**Fig. 1E-G, Fig. S1B-G**). We compared the Probound predictions (**Fig. 1E**) with *in vitro* gel shift experiments and observed a linear dependence over the range of affinities we tested (**Fig. 1F**, **Fig. SS2**) indicating agreement between the Probound binding model and relative U2AF binding affinity measured by experiment. Surprisingly, we find that U2AF occupies highly degenerate binding sites *in vivo* and that annotated 3’SSs contain only marginally stronger U2AF binding sites than sites at other locations, as shown for example in *RAB7A* (**Fig. 1G**). Therefore, we sought to address how fractional differences in equilibrium binding affinity could result in observed splicing outcomes by examining U2AF binding kinetics to RNA.

We implemented a fluorescence assay to determine the kinetics underlying the U2AF binding affinity landscape. This approach, called **PIFE** (*21*) (Protein Induced Fluorescence Enhancement), describes a phenomenon where protein binding to a site immediately neighboring a Cy3 fluorophore results in an increase in fluorescence emission. When titrating U2AF against the well characterized Adenovirus Major Late Promoter (‘AdML’) 3’SS containing a Cy3 after the 3’SS (***see methods***), we measure a 1.5-fold increase in fluorescence emission (**Fig. 1H)**. Furthermore, the data can be easily fit to a non-cooperative Hill Equation and the resulting K_D_ closely matches the EMSA measurement K_D PIFE_ = 27 ± 7 nM (**Fig. 1I**), indicating that this approach indeed detects U2AF binding. We extended this assay to the single-molecule level to measure binding kinetics where Cy3-labeled surface tethered RNAs are visualized by Total Internal Reflection Fluorescence microscopy (**TIRF**) (**Fig. 1J**). Time traces were fit to a two-state HMM model where high fluorescence corresponds to RNA in complex with U2AF and low fluorescence corresponds unoccupied RNA **(Fig. 1K)**. We measured U2AF binding to two 3’SSs, AdML and a similar 3’SS but with three uridine mutations in the py-tract (CCC**<u>C</u>**<u>U**A**UU**C**UU</u>CC) (**Fig. 1L, Fig. S3A-B, *see methods for full sequence***). In the single-molecule assay, we measure essentially the same overall association rate (**Fig. 1M**) and a 1.5 ± 0.2-fold slower dissociation rate from the AdML sequence (**Fig. 1N**, **Fig. S3C-G**). Therefore, the reduction in binding affinity to the mutant sequence is due exclusively to faster dissociation from RNA. From this measurement, we conclude that association rate is not sensitive to sequence and that binding affinity differences are due to changes in the U2AF dissociation rate from RNA. Taken together, we find that U2AF interacts for about 15 seconds to the ‘ideal’ AdML U2AF 3’SS *in vitro* **(Fig. 1N)**. Considering a median strength U2AF site in human pre-mRNA is ∼100-fold weaker (**Fig. 1O)**, our analysis predicts U2AF binds on the sub-second time scale to bona fide 3’SSs *in vivo*. It is not known how these transient interactions lead to correct 3’ SS selection against the background of highly degenerate sites in pre-mRNA.

### SIMULTANEOUS OBSERVATION OF U2AF BINDING AND PRE-mRNA SPLICING IN LIVING CELLS

We therefore speculated that while transcriptome-wide U2AF binding affinity imparts an initial subtle bias towards bona fide 3’SSs, most specificity is accomplished through cooperative interactions as the spliceosome assembles (*22*). To test this idea, we sought to simultaneously image splicing factor binding and intron removal on a single pre-mRNA in living cells. To begin, we developed an approach to visualize formation of the A complex on endogenous pre-mRNAs where all regulatory elements are present. We used CRISPR to integrate a HALO-tag on the C-terminus of U2AF1 **(Fig. 2A**) or U2AF2 (**Fig. S4**), allowing us to image U2AF binding in live cells at endogenous concentrations. Based on *in vitro* studies, we expected most U2AF-RNA interactions to occur for less than one second (**Fig. 1O**). Therefore, we initially tracked single U2AF1-HALO proteins at an acquisition rate of 100 Hz to resolve both bound and unbound events (*referred to as ‘fast SMT’ herein*) (**Fig. 2B-C, Fig. S4**). When analyzing binding events that start and end within the short duration of the video, we do indeed observe a population of transient binding events on the sub-second time scale (**Fig. 2D**). However, we also observe a small population of stable binding events that persist for long periods and, when tracking U2AF-HALO over much longer time scales (20 minutes, 0.33 Hz, *referred to as* “*slow SMT’* herein), we observe a broad dwell time distribution that appears linear on a log-log plot with the longest binding events occurring for over 100 seconds (**Fig. 2E, Fig. S4D for kymographs**). Considering our finding that 3’SSs predominantly contain degenerate U2AF binding sites (**Fig. 1O**), these long dwell times, which extend past the timescales of U2AF binding high-affinity sequences *in vitro* (**Fig. 2E**), likely reflect U2AF binding RNA as a member of a larger, stabilizing complex.

**Figure 2:**
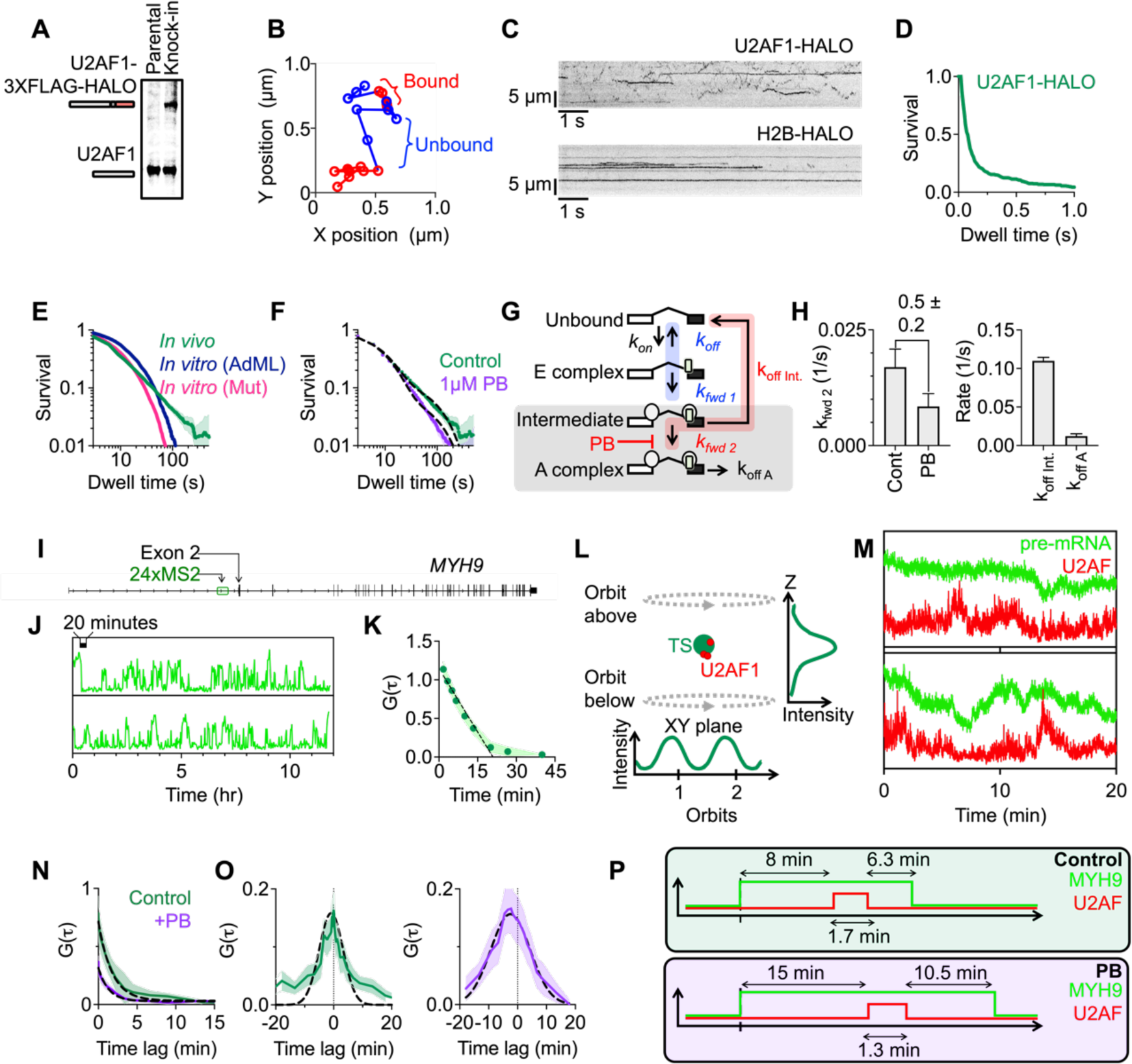
Single-molecule visualization of spliceosome assembly reveals stable U2AF binding through multiple assembly intermediates. A) Western blot of HBEC line containing one copy of U2AF1-3XFLAG-HALO, subsequently referred to as U2AF1-HALO. B) Fast tracking of U2AF1-HALO diffusion throughout the nucleus imaged with Highly Inclined and Laminated Optical Sheet (HILO) illumination at 100 Hz over 10 seconds. Bound and unbound states were inferred using a 2 state HMM model. We analyzed tracks that show complete binding events (i.e. bound event flanked by two unbound events) (n=2 replicates). C) Kymographs comparing U2AF1-HALO and H2B-HALO diffusion in fast tracking assays. D) Dwell times of complete binding events measured in fast tracking assays. E) Slow tracking U2AF1-HALO diffusion throughout the nucleus at 0.33 Hz over 20 minutes (n=3 replicates, shaded region represents 95% CI). For comparison, *in vitro* U2AF dwell time distributions from Fig. 1N are also included. F) Treating cells with pladienolide B (PB) results in a reduction in U2AF1-HALO dwell times (n=3 replicates). Shaded region represents 95% CI. G) Interpreting slow SMT data which detects U2AF dissociation from spliceosome complexes with a splice site selection model (intermediate and A complex states highlighted in gray box). Data is fit so that k_off Int._ and k_off A_ are the same for both control and PB datasets. Blue highlighted rates represent K_f_, the ratio of spliceosome forward progression with respect to U2AF dissociation at the E complex (K_f_ = k_fwd 1_/ k_off_). Red highlighted rates represent K_s,_ the ratio of spliceosome forward progression with respect to U2AF dissociation at the intermediate state (K_s_ = k_fwd 2_/ k_off Int._). H) Model fitting reveals pladienolide B reduces k_fwd 2_, the transition rate from intermediate to A complex by 2-fold. Additionally, our fits indicate U2AF remains bound in the intermediate state for 9.1 ± 0.4 seconds and the A complex for 82 ± 21 seconds. I) We used CRISPR to integrate a HALO tag on the C-terminus of U2AF1 in a cell line containing 24 MS2 stem loops integrated about 4kb upstream of the first 3’SS in MYH9. These cells also contain GFP-MS2 coat protein (MCP) so that upon transcription of stem loops, MCP binds and enables detection of RNA. J) Overnight confocal imaging of MYH9 transcription sites reveals both transcription (transition to high fluorescent state) and splicing (transition back to low fluorescent state). K) Fluctuations were analyzed by correlation analysis and indicates a characteristic time of 19 ± 2 min for transcription and splicing. Error bars represent one standard deviation. L) 3D orbital tracking: a laser orbits above and below the transcription site. By monitoring the intensity in the XY plane as well as in Z, the particle is tracked in three dimensions. We tracked MYH9-GFP while monitoring U2AF diffusion in and out of the confocal volume in another channel. M) Example traces of MYH9 splicing and U2AF binding. Imaging was performed at 4 Hz over 20 minutes. Splicing manifests as a sudden decrease in GFP signal. N) Autocorrelation of time traces in U2AF channel corresponds to dwell times of τ_control_ = 1.7 ± 0.1 min, τ_PB_ : 1.3 ± 0.1 min. Error bars represent one standard deviation. O) U2AFèMYH9 cross-correlations are centered around 0 indicating overlap between MYH9 and U2AF events. Additionally, the width of the cross-correlation curves represent the time to splice (τ_splice control_ = 16 ± 1 min, τ_splice PB_ = 27 ± 1 min). Error bars represent one standard deviation. P) Summary of findings from SMT and OT experiments. U2AF binds for about 2 minutes to a pre-mRNA undergoing the splicing reaction. While U2AF dwell times are shorter after treatment with PB, overall time to splice increases.

To test whether these long dwell times represent splicing complexes containing U2AF, we treated our cells with 1 uM pladienolide B (PB), a splicing inhibitor that blocks formation of the A complex (*23*, *24*). Indeed, we observe a shift in the dwell time distribution towards shorter dwell times (**Fig. 2F**). However, even after PB treatment, the time scales of binding are still longer than possible for an isolated U2AF-RNA interaction. Therefore, in contrast to previous measurements (**Fig. 1 & 2B-D**), both experiments (Control and PB) likely reflect U2AF dissociation while in complex with the spliceosome, with the longest dwell times corresponding to U2AF dissociation after successful splicing initiation and shorter time scales representing dissociation from an earlier intermediate state which becomes enriched after PB treatment. We modeled this relationship as shown in **Fig. 2G**. This general model, motivated by previous biochemical and structural data, is intended to encompass a wide range of possible behavior including reversible, irreversible and proofreading steps. We obtain the following equation for the survival curve with three time constants, which will be constrained by time-resolved measurements of U2AF binding in cells. (**Fig. S4G for master equations)**:

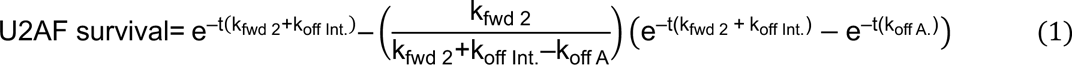

We assume that PB affects only k_fwd 2_, the rate of forward progression from the intermediate state to the A complex. Therefore, both datasets were fit simultaneously so that k_off Int._ and k_off A_ were shared. With these conditions, we find that PB treatment leads to a 2-fold reduction in k_fwd 2_. Furthermore, the fit values for k_off int._ and k_off A_ correspond to lifetimes of 9.1 ± 0.4 seconds and 82 ± 21 seconds for the Intermediate and A states, respectively (**Figs. 2F black line, 2H**). While the slow SMT assay cannot resolve the initial transient binding involved in the regulation of splice site selection at the E complex, it does resolve the regulation of splice site selection around the intermediate state. Importantly, these experiments establish that U2AF dissociation out of the intermediate state is much faster than spliceosome forward progression, indicating that the majority of U2AF molecules in the intermediate state dissociate to terminate spliceosome assembly.

The prediction from this kinetic analysis is that U2AF undergoes repeated binding events throughout the splice site selection process resulting in long interactions with the binding site only during a successful splicing reaction. Thus, we aimed to simultaneously image splicing and U2AF binding. We created a cell line containing, in addition to U2AF1-HALO, 24xMS2 stem loops integrated about 4 kb upstream of the first 3’SS in *MYH9* (**Fig. 2I**). When transcribed, GFP-MS2 coat proteins bind the stem loops enabling visualization of the RNA (*6*, *25*, *26*). We chose this pre-mRNA for the following reasons: (i) while U2AF binds pervasively throughout introns, we observe no extra U2AF binding sites between the MS2 integration and the 3’SS, and (ii) single-molecule FISH reveals that *MYH9* is transcribed in most cells but generally contains fewer than two RNAs per transcription site (**Fig. S4H**). Indeed, when imaging MYH9 splicing kinetics by confocal imaging, we observe two states: the transition from low to high fluorescence represents transcription and the transition from high to low fluorescence represents splicing (**Fig. 2J**). We analyzed the fluorescent time traces by correlation analysis in the same way as previous studies (*27*, *28*) and measured a ‘time to splice’ (**ι−_splice_**) of 19 ± 2 min, which corresponds to synthesis of the stem loops, elongation to the 3’SS, and subsequent intron removal (**Fig 2K**), consistent with previous measurements (*6*, *29*).

We overcame the technical challenge of imaging both protein binding (which occurs for seconds) and pre-mRNA splicing (which occurs on the ∼15 minute time scale) with orbital tracking spectroscopy. The *MYH9* transcription site is tracked in three dimensions using a laser that orbits above and below the transcription site (*30*) (**Fig. 2L**). While tracking *MYH9*, we also monitor the corresponding signal in the U2AF channel. While most measurements begin in the ‘high’ state when the MS2 signal is visible and end after splicing (**Fig. 2M**, top), some tracks also show the full cycle of transcription and splicing **(Fig. 2M**, bottom). Autocorrelation of the U2AF signal reveals the timescales of U2AF binding in a productive splicing reaction and when splicing is stalled by PB (Control: 102 ± 6 s, PB: 78 ± 6 s) **(Fig. 2N**). These dwell times are within measurement error of the time required to dissociate from the A complex in the slow SMT assays (**Fig. 2H**). Furthermore, the U2AF-RNA cross-correlation curves can be approximated with gaussian distributions where the width represents ι−_splice_ (Control = 16 ± 1 min, PB = 27 ± 1 min) and the center position represents the time delay from the middle of the U2AF binding event to the middle of the *MYH9* splicing event (Control = -0.9 ± 0.2 min, PB = -2.5 ± 0.1 min) **(Fig. 2O**). The broadening of the cross-correlation curve after PB treatment represents slower overall splicing in this condition. Taken together, this data reveals the time scales of U2AF binding during *MYH9* splicing **(Fig. 2P**) and further supports a model where U2AF binds stably to actively spliced pre-mRNAs after progressing through transient initial states.

### A KINETIC MODEL FOR SPLICE SITE SELECTION

Our *in vitro* and *in vivo* assays indicate extensive use of moderate and low affinity U2AF sites where accurate splice site selection must occur despite sub-second dwell time differences between functional and non-functional sites. The core element of this model is ATP-dependent irreversible forward progression (k_fwd 1_ and k_fwd 2_) in competition with U2AF dissociation (k_off_ and k_off int_) (**Fig. 2G**) where we assume that both dissociation steps depend on underlying sequence. Repeating this scheme twice (around the E complex and Intermediate state) is a hallmark of kinetic proofreading and has the potential to greatly improve the fidelity of splice site selection at bona fide splice sites (*31*).

To test the ability of this model to predict *in vivo* splicing decisions, we related alternative splicing patterns with the underlying U2AF equilibrium binding affinity by performing RNA sequencing in Human Bronchial Epithelial Cells (HBECs) followed by SUPPA2 alternative splicing analysis (*32*). We were particularly interested in alternative 3’SS selection where two closely spaced 3’SSs compete for U2AF. Applying the kinetic model from **Fig. 2G** to alternative 3’SS selection (**Fig. 3A, 3B**), we get the following equation for Percent Spliced In (**PSI**) (*see **Fig. S5A** for master equation)*:

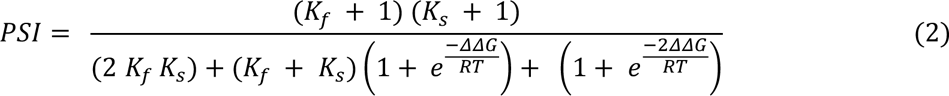

**Figure 3:**
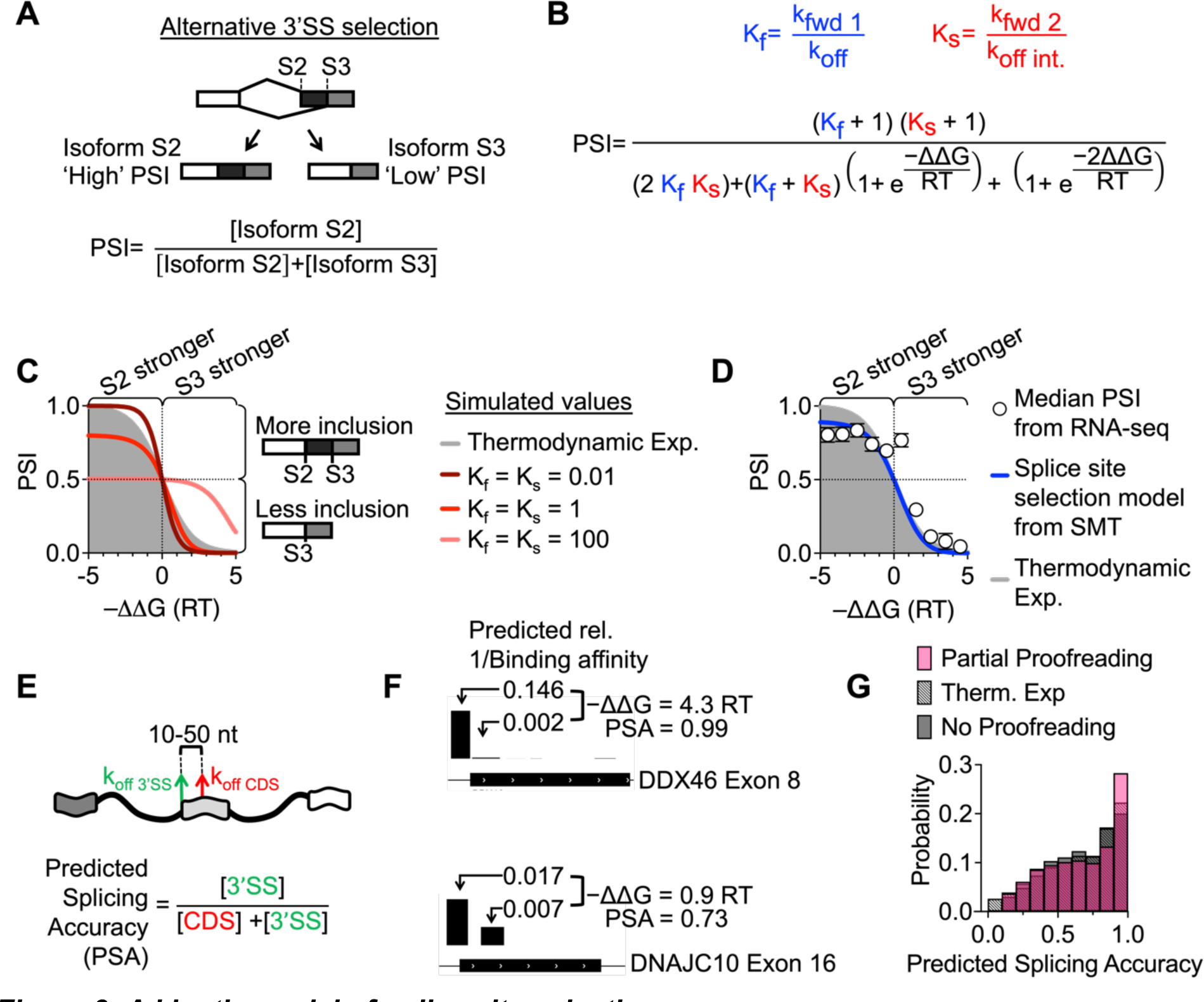
A kinetic model of splice site selection A) Alternative 3’SS selection. U2AF may bind to initiate spliceosome assembly at two competing 3’SSs, S2 and S3. B) Applying the kinetic model developed in figure 2 to the process of alternative 3’SS selection. This equation relates relative U2AF dissociation rate (−ΔΔG = RT ln(k_off 3_/k_off 2_)) to percent spliced in (PSI). While the Probound binding model in Fig. 1 initially reports relative binding affinities instead of dissociation rates, we assume only dissociation rates (and not association rates) depend on sequence and, therefore, use k_off_ instead of K_D_ in our −ΔΔG calculation. C) The predicted relationship between −ΔΔG and PSI described by equation 2. The gray plot shows the expected relationship between PSI and ΔΔG in the scenario where splice site selection depends entirely on the relative equilibrium binding affinity between sites S2 and S3. The maroon curve represents the ‘full proofreading’ scenario where U2AF reads the 3’SS twice and both K_f_ and K_s_ << 1. Increasing K_f_ and K_s_ reduces the role of underlying U2AF binding affinity in splice site choice. For example, when K_f_ and K_s_ = 100 (salmon curve), PSI asymptotes at 0.5 at negative −ΔΔG. D) We performed RNA sequencing and quantified alternative splicing in HBECs. PSI values were binned by −ΔΔG, the free energy of U2AF binding the S3 site relative to the S2 site. White circles represent median PSI of each bin. Error bars represent the average standard deviation of PSI for each alternative 3’SS (n = 2 replicates). The blue curve represents the predicted relationship between PSI and −ΔΔG based on slow SMT measurements (K_s_ = 0.15). K_F_ can be fixed to any value above 10. E) Determining predicted splicing accuracy (PSA) by applying equation 2 for U2AF binding affinity at 3’SSs relative to sites in downstream coding regions (10-50 nt downstream). F) Comparing predicted U2AF binding affinity at 3’SSs with respect to downstream sites in coding regions. The height of each bar corresponds to 1 / relative binding affinity. We used the measured rates from the SMT assays to calculate PSA. G) The distribution of predicted splicing accuracy (PSA) for a non-proofreading ‘bind-once’ model (gray, K_s_ = 0.15), thermodynamic expectation (patterned), and partial proofreading model (pink, K_f_ = 10 and K_s_ = 0.15). The partial proofreading model predicts the highest splicing accuracy. U2AF binding locations come from PAR-CLIP measurements and we only measured pairs of sites where predicted relative binding affinity for both sites is more than 0.1% of the highest affinity sequence.

Here, we normalize the kinetic quantities as unitless parameters K_f_ = k_fwd 1_ / k_off_ and K_s_ = k_fwd 2_ / k_off Int_, which we measured with the SMT assays. ΔΔG reflects the difference in Gibb’s free energy of U2AF binding to two potential 3’SSs (**Fig. 3A**), measured directly with RBNS and related to the off rates of U2AF from the two potential sites by 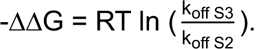. The parameters K_f_ and K_s_ reflect the importance of U2AF binding site strength in determining splicing outcome. If binding sites are very weak (K_f_ and K_s_ << 1), forward progression is rare, and the probability of achieving a specific mRNA isoform is highly dependent on the -ýýG of U2AF binding (**Fig. 3C, maroon curve**). Additionally, compared to a model where PSI depends *only* on relative U2AF binding affinity at a single step between two potential 3’SSs (**Fig. 3C, gray,** ‘thermodynamic expectation’), this model is more sensitive to underlying binding affinity by emphasizing the proofreading branch, as conveyed by the steepness of curves when K_f_ and K_s_ are sufficiently low. We call this regime ‘full proofreading.’ In contrast, if forward progression is much faster than U2AF dissociation (K_f_ and K_s_ >> 1), PSI ∼ ½, and there will be no specificity based on binding site strength, because *any* binding event leads to forward progression (**Fig. 3C, salmon curve**). Importantly, this scenario highlights how high affinity binding (i.e. small k_off_) can be unfavorable in regulating alternative 3’SS selection. We call this regime ‘no proofreading.’ Finally, the regime of ‘partial proofreading’ (K_f_, K_s_ ∼ 1) displays both better and worse discrimination than the thermodynamic expectation, depending on ýýG. Note, that both ‘no’ and ‘partial’ proofreading models emphasize irreversible forward progression, leading to an asymptote at values below PSI=1 at large negative −ΔΔG which corresponds to the use of weak annotated 3’ SS splice sites that would never be used at equilibrium.

A clear sigmoidal relationship emerges when calculating the median PSI of groups of alternative 3’SSs binned by -ýýG as determined with RBNS (**Fig. 3D**, white circles), even though there is variation at the level of individual transcripts **(Fig. S5B).** Taken in context with our earlier binding studies, this result emphasizes the surprisingly strong influence of moderate and weak U2AF binding sites in determining alternative splicing patterns. The overall shape indicates both equilibrium and non-equilibrium elements and suggests that alternative 3’SS selection does not operate in the ‘full proofreading’ regime where both K_f_ and K_s_ << 1 and specificity is optimized. Rather, using the ‘partial proofreading’ model with K_s_ = 0.15 measured in the slow SMT assay (**Fig. 2H**), we find that the PSI dependence is best fit with K_f_ ≥ 10 (**Fig. 3D**, blue line) which indicates U2AF binding leads to rapid initial progression toward the A complex with 3’SS selection occurring after initial binding during subsequent assembly steps.

We next consider the scenario where U2AF must identify the correct annotated 3’SS with respect to nearby downstream sites, for example in coding sequences. This selection cannot be readily identified by RNA-seq due to quality control of mis-spliced transcripts. As such, we computed the predicted splicing accuracy (PSA) (**Fig. 3E**) -- analogous to PSI -- and compared this quantity to the thermodynamic expectation where PSA is only due to equilibrium binding affinity. PSA was computed by using the RBNS ýýG for a U2AF PAR-CLIP peak at the 3’ ss vs. the nearest downstream exon site, as shown for example in **Fig. 3F**. Importantly, the thermodynamic expectation describes a scenario where spliceosome forward progression is infinitely slow so that equilibrium can be reached. In contrast, our partial proofreading model achieves a higher predicted splicing accuracy despite allowing for spliceosome forward progression (**Fig. 3G**). In fact, the most common prediction is that more than 90% of spliceosomes correctly assemble at annotated 3’SS and not at nearby downstream sites in coding sequence.

### DDX42 PROMOTES SPLICING SPECIFICITY BY ACCELERATING U2AF DISSOCIATION

We sought to identify the molecular players in this partial proofreading scheme. Specifically, we aimed to identify proteins that interact with U2AF during spliceosome assembly and to apply our single-molecule approach to reveal how these proteins influence spliceosome progression. To identify proteins that interact with U2AF, we used CRISPR to insert a FLAG-tag on the C-terminus of U2AF1 and performed Immunoprecipitation-Mass Spectrometry (IP-MS) on HBEC nuclear extracts. Surprisingly, the most enriched splicing factor was DDX42, a relatively uncharacterized helicase that interacts with SF3B1 during the spliceosome assembly process (*33–35*) (**Fig. 4A- C**).

**Figure 4.**
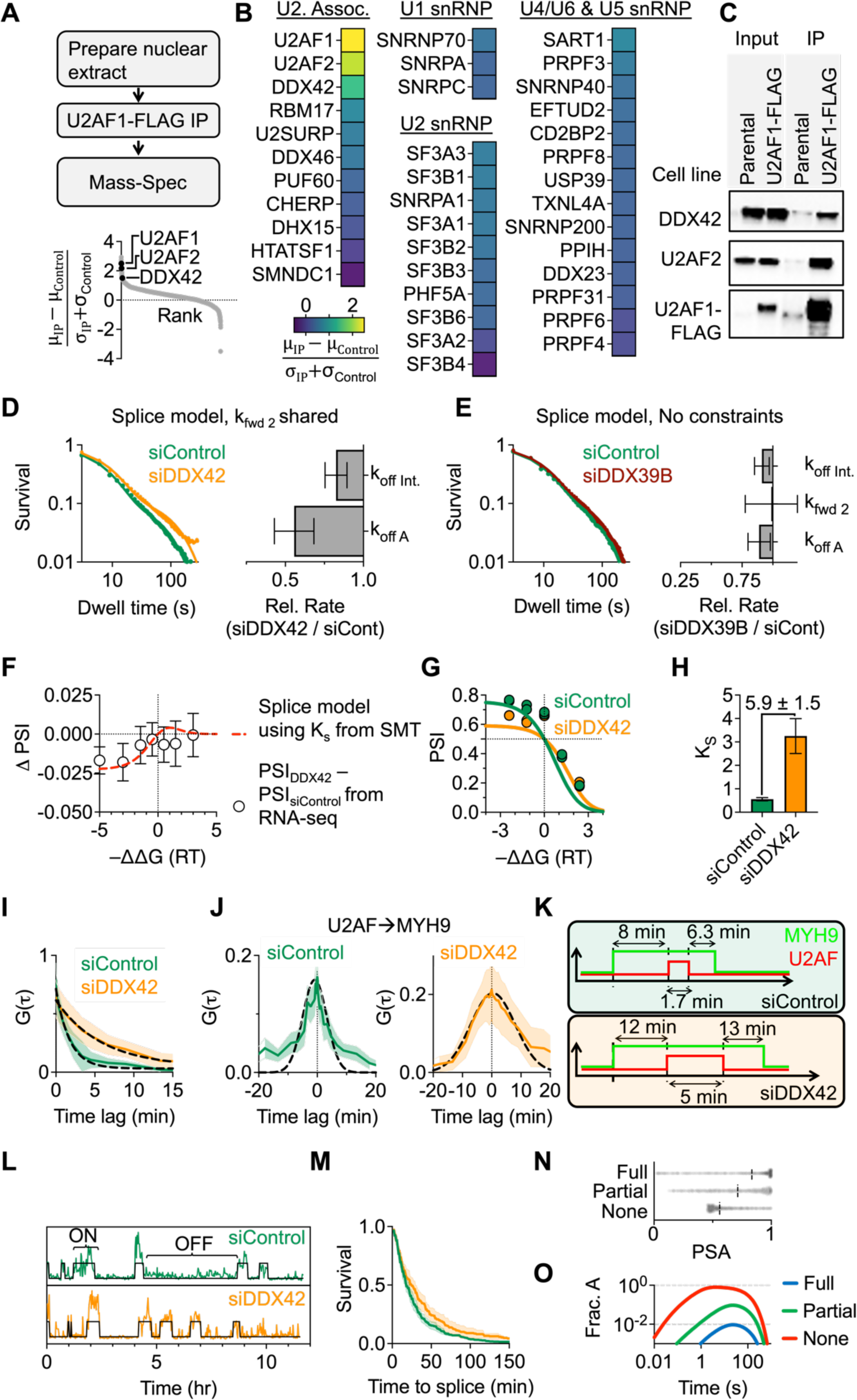
DDX42 accelerates U2AF dissociation from intermediate state and A complex A) Top: Workflow for U2AF1-FLAG IP-MS from HBEC nuclear extract. Bottom: Ranking identified proteins by signal-to-noise (where control is parental HBEC cell line). Other than U2AF1 and U2AF2, DDX42 is the most enriched splicing factor (n = 3 replicates). B) Plotting enrichment of proteins associated with U1, U2, and the tri-snRNP from our U2AF1 IP-MS assay. The protein components of each snRNP were determined using the spliceosome database (*38*). C) Western blot after U2AF1-FLAG IP confirms U2AF1-DDX42 interaction. D) Comparing U2AF1 dwell time distributions in the slow SMT assay after electroporation with DDX42 or Control siRNA. Fitting to the kinetic model from Fig. 2G reveals a reduction in k_off Int.,_ and k_off A_ . Error bars in bar chart represent standard error of the mean from 3 replicates. E) Measuring U2AF dwell times after knockdown of DDX39B (n = 1 replicate). Error bars represent standard deviation of 1000 bootstrapping iterations. F) Predicting the change in PSI (PSI_siDDX42_ – PSI_siControl_) after DDX42 knockdown based on the change in K_S_ measured in slow SMT. Red dashed lines represent predicted PSI, white circles represent mean ΔPSI for binned −ΔΔG values from RNA sequencing. This analysis was performed on alternative 3’SSs where U2AF affinity to both sites was within 50-fold of the ideal site (6994/24112 alternative 3’SSs). G) The relationship between PSI and ΔΔG for the 854 3’SSs that exhibit significant splicing changes after DDX42 knockdown (P<0.05). H) For this subset of alternative 3’SSs, DDX42 knockdown increases K_S_ by 5.9 ± 1.5-fold. Error bars represent one standard deviation. I) Autocorrelation of the U2AF time traces from the orbital tracking measurements after DDX42 knockdown show that, on a pre-mRNA undergoing the splicing reaction, U2AF dwell times are longer (τ_bound U2AF siDDX42_ = 5.8 ± 0.2 min, τ_bound U2AF siCont_ = 1.7 ± 0.1 min). Error bars represent one standard deviation. J) U2AFèMYH9 cross-correlation broadens after DDX42 knockdown, indicating slower overall splicing (τ_splice U2AF siDDX42_ = 30 ± 1 min, τ_splice U2AF siCont_ = 16 ± 1 min). Error bars represent one standard deviation. K) Interpretation of SMT and orbital tracking data: DDX42 knockdown prolongs both U2AF binding and MYH9 splicing. L) Example time traces of gene trap cell line showing transcription and splicing events. M) Measuring changes splicing kinetics globally with the gene trap cell line τ_1/2 siControl_ = 18 min, τ_1/2 siDDX42_ = 26 min. Shaded regions represent one SEM (n=4 replicates for siControl, n=2 replicates for siDDX42). N) Comparing predicted splicing accuracy (as in figure 3E-G) for 3’SSs in competition with binding sites in nearby downstream coding regions for three regimes of the proofreading model, full (K_f_ = 0.1, K_s_ = 0.15), partial (K_f_ = 10, K_s_ = 0.15), and none (K_f_ = 10, K_s_ = 10). O) Comparing the fraction of bound U2AF molecules in the A complex for the three scenarios introduced in N. While figure N establishes that full proofreading is the most accurate, it is also very slow, at best, only ∼1% of bound U2AF molecules are in the A complex at a specific time.

We performed SMT assays after DDX42 knockdown to determine if this protein influences U2AF binding kinetics and subsequent spliceosome assembly (**Fig. S6A**). Fast SMT assays indicate DDX42 knockdown has no impact on the time scales of transient sub-second U2AF-RNA interactions (**Fig. S6B-C**). However, the slow SMT assay, which only detects stable U2AF binding, including U2AF in complex with the spliceosome, shows slower U2AF dissociation across time scales ranging from 10-300 seconds after DDX42 knockdown (**Fig. 4D**, **Fig. S6D**) but not after depletion of another helicase believed to be involved in formation of the early spliceosome, DDX39B (*36*) (**Fig. 4E, Fig. S6E**) These results suggest that DDX42 interacts with U2AF during the process of spliceosome assembly, not during initial binding site sampling. We fit the slow SMT data to the same splice model as in **Fig 2G (Eqn. 1).** Model testing indicates the data fits best when k_fwd 2_ is shared and the two dissociation rates are free to fluctuate (**Fig. 4D, Fig. S6F-G**). Therefore, in contrast to pladienolide B which slows spliceosome forward progression, the absence of DDX42 seems to slow U2AF dissociation. Our fit indicates a subtle, 18 ± 7% decrease in the U2AF dissociation rate from the Intermediate state (k_off int_) and a larger 44 ± 13% decrease in U2AF dissociation from the A complex (k_off A_).

Our kinetic model of alternative splice site selection (**Eqn. 2**) predicts that the ∼20% change in K_s_ measured by slow SMT upon DDX42 knockdown will result in a small global reduction in PSI for alternative 3’SS selection, especially in the negative -ýýG regime where U2AF binds the upstream 3’SS site more tightly (**Fig. 4F**, dashed red curve). To test this, we performed RNA-seq after DDX42 knockdown and applied our alternative splicing analysis. Indeed, when binning ýPSI values for alternative 3’SSs after DDX42 knockdown into groups of similar relative U2AF binding affinity (**Fig. 4F**, open circles), we observe excellent agreement with our SMT data, both in terms of the magnitude of splicing changes and in terms of the trend of the relationship. Importantly, this result establishes that a subtle change in U2AF dissociation from the intermediate state is sufficient to affect alternative splice site selection globally. In addition to this global analysis, we also identified 3’SSs that show significant changes in alternative 3’SS selection (P<0.05). While only 854/12514 (7%) of alternative 3’SSs meet this criteria, we measure nearly a 6-fold increase in K_s_ (5.9 ± 1.5-fold) for this subset of alternative 3’SSs (**Fig. 4G-H**). While the magnitude of this change is much greater than predicted from the SMT assays, both experiments are consistent with slower U2AF dissociation from the intermediate state after DDX42 knockdown. Therefore, we conclude that DDX42 evicts U2AF from pre-mRNA in the partial proofreading scheme.

Finally, we interrogated the role of DDX42 in spliceosome progression using additional live-cell assays. In the orbital tracking assay, we observed a substantially slower U2AF dissociation after DDX42 knockdown (τ_U2AF control_ = 1.7±0.1 min, τ_U2AF siDDX42_ = 5.8 ± 0.2 min) (**Fig. 4I**) consistent with the global slowdown of U2AF dissociation from the A complex after DDX42 knockdown observed in slow SMT. We also observed a broader cross-correlation distribution (τ_U2AF control_ = 16 ± 1 min, τ_U2AF siDDX42_ = 30 ± 1 min) indicative of slower overall splicing (**Fig. 4J**, **Fig. 4K**). Lastly, to determine the pervasiveness of this effect, we measured splicing dynamics globally using a polyclonal gene trap where MS2 stem loops are inserted into many introns. This cell population contains MS2 integrations in 728 unique introns in HBECs (*6*), allowing us to generate a ‘dynamic metagene’ analysis of intron removal kinetics. We see that DDX42 knockdown results in a global slowing of pre-mRNA splicing (**Fig. 4L-M**). In summary, we conclude that knockdown of DDX42 results in slower U2AF dissociation (predominantly from the A complex in our model), leading to altered kinetic discrimination of splice sites, slower overall spliceosome progression, and consequently changes in alternative splicing.

## DISCUSSION

Here we illuminate the dynamic operating principles of the early spliceosome. At the heart of this description is a basic kinetic module consisting of equilibrium association of an RNA binding protein, followed by an irreversible, energy-consuming transition to a complex that can be proofread and disassembled as needed (e.g. **Fig. 2G**). We find that 3’SSs generally contain weak U2AF binding sites: U2AF initially binds a potential 3’SS for less than a second. Despite these weak sites, the spliceosome is remarkably sensitive to subtle differences in U2AF binding affinity. U2AF recognition has thus adopted a strategy not on discrimination at the initial bound state but on rapid progression to an intermediate (K_f_>>1) which is subject to proofreading. In contrast, progression from the intermediate to the A complex is inefficient (K_s_<<1), which leads to repeated sampling of the binding site before progression. This strategy may be successful because U2AF is highly expressed in the nucleus which could enable very fast association to pre-mRNAs (*37*). Furthermore, while operating in ‘full’ proofreading mode (K_f_<<1, K_s_<<1) would achieve the best binding site discrimination (**Fig. 4N**), it comes at the cost of inefficient progression (**Fig. 4O**). Therefore, the partial proofreading model seems to both (i) discriminate based on binding affinity better than possible for the simple thermodynamic model while (ii) enabling efficient forward spliceosome progression. In this partial proofreading scheme, we identify a role for DDX42 in accelerating U2AF dissociation. In this way, DDX42 increases spliceosome specificity to the underlying U2AF binding site.

## AKNOWLEDGEMENTS

We thank members of the Larson laboratory for helpful discussions. We thank the many support staff at the NIH core facilities who maintained the instruments and provided helpful advice when conducting experiments, including Dave Ball (NCI Optical Microscopy Core), Jiji Chen (Advanced Imaging and Microscopy Reseource), Di Wu (NHLBI Biophysics Core), Gianluca Pegoraro and Adib Keikhosravi (NCI High-Throughput Imaging Facility), Madeline Wong (CCR genomics core), the Frederick Protein Expression Laboratory, the CCR Protein Characterization Laboratory, the CCR sequencing facility, and the NIH High-Peformance Computing Group (Biowulf). This research was funded by the Intramural Research Program of the NIH.

## METHODS

### Expression and purification of full length U2AF heterodimer

DNA fragments coding for 3X-FLAG-U2AF1 and 6X-HIS-U2AF2 were commercially synthesized (IDT) and cloned downstream of the p10 and polyhedron promoters respectively in the pFASTBAC baculovirus expression vector. Sf9 cells were transduced with baculovirus packaged with the expression vectors in the NIH protein expression laboratory. Cells were stored as pellets at -80C until ready to purify. Cell pellets were thawed on ice and resuspended in Buffer A (50 mM tris pH 7.5, 5 mM imidazole, 10% glycerol, protease inhibitor tablet (Pierce Protease Inhibitor Tablets, EDTA-free)). After thawing, cells were lysed by sonication with 15 cycles (15 s on/10 s off) using the Bioruptor Plus (Diagenode) at 4C. The first 5 cycles were set to ‘high’ power and the last 10 cycles were set to ‘low’ power. After lysis, Cells were clarified by centrifugation at 23000xg for 20 min. Supernatent was manually loaded on to 5 mL His-trap HP IMAC column very slowly and then installed onto a BioRad NGC FPLC. We performed 30 mL wash with buffer A to remove unbound protein followed by a 60 mL wash with Buffer B (50 mM tris pH 7.5, 60 mM imidazole, 10% glycerol). We observe a very large elution peak corresponding to U2AF2 (without U2AF1) at this imidazole concentration. The full U2AF heterodimer eluted when stepping into buffer C (50 mM Tris pH 7.5, 340 mM imidazole, 10% glycerol). To ensure each U2AF2 molecule was in complex with U2AF1, we finished the purification by U2AF-IP with anti-FLAG M2 magnetic beads (Sigma Aldrich). Before mixing with U2AF, beads were washed 4x in Buffer D (10 mM tris pH 7.5, 500 mM nacl, 0.1% tween20, 0.5 mM DTT, protease inhibitor tablet. U2AF was eluted with 25 mM Tris pH 7.5, 100 mM NaCl, 200 ug/ml 3xflag, 1mM DTT. Protein concentration and purity was assessed by SDS PAGE. Protein was stored at -80C after adding glycerol to 10%.

### Electrophoresis Mobility Shift Assays (EMSAs)

5 nM RNA was mixed with 0-120 nM U2AF in binding buffer (25 mM Tris pH 7.0, 150 mM NaCl, 0.5 mM DTT, 10% glycerol, 0.05% Tween 20, 0.02 ug/ul tRNA) and incubated at 4C for 15 minutes. Binding was resolved in a native PAGE gel (1XTBE, 8% polyacrylamide (29:1 National Diagnostics), 0.1% APS, 0.1%TEMED, 10% Glycerol) run at 75 volts for 2 hours at 4C. Gel was pre-run for 1 hour before loading.

We measured binding to the following sequences (changes to mutant sequence in bold). The red region is a DNA sequence used for tethering RNAs to microscope slide in single-molecule PIFE assays.

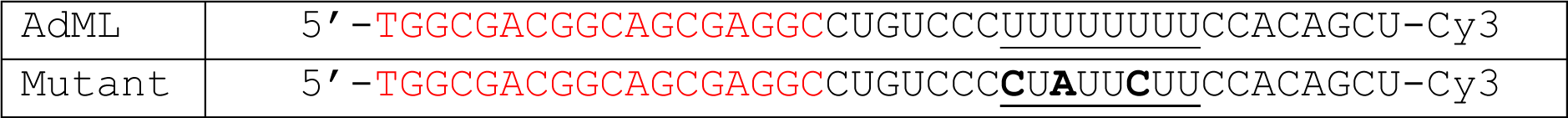

### RNA Bind-N-Seq (RBNS)

Our protocol for RNA Bind-N-Seq closely follows the protocol developed by the Burge Lab (*39*). The random RNA library was prepared by annealing the T7 oligo to the RBNS template, which is a single-stranded DNA oligo from IDT (sequences are listed in table below). We then performed *in vitro* transcription for 4 hours using the NEB HiScribe T7 RNA synthesis kit. After RNA synthesis, we degraded the DNA template by DNaseI digestion for 15 minutes. RNA was purified using the NEB Monarch RNA cleanup kit and stored at -80C. We confirmed that RNA was the correct length by tapestation analysis (Agilent).

U2AF (0 nM, 20 nM, 80 nM, or 320 nM) was incubated with 500 nM of random RNA library in U2AF binding buffer without tRNA (25 mM Tris pH 7.0, 150 mM NaCl, 0.5 mM DTT, 10% glycerol, 0.05% Tween 20) at 4C for 30 minutes with constant mixing on a rotisserie tube rotator. We performed each binding reaction in duplicate. Final volume was 250 uL per reaction. RNA in complex with U2AF was isolated by U2AF1-FLAG IP using anti-FLAG M2 magnetic beads (Sigma Aldrich). Beads were washed 4x in RBNS wash buffer (25 mM Tris pH 7.5, 150 mM KCl, 60 ug/mL BSA, 0.01% Tween 20) and then resuspended in 200 uL of binding buffer. We added 20 uL of washed beads to each RBNS reaction and incubated for 30 minutes at 4C. Bound RNA was pelleted with magnetic beads and incubated in 1000 uL RBNS wash buffer for 15 minutes at 4C. Next, the RNA was pelleted and resuspended in elute buffer (25 mM HEPES pH 7.3, 100 mM NaCl, 200 ug/ml 3XFLAG (sigma), 1 mM DTT). U2AF was removed from RNA by phenol-chloroform extraction and ethanol precipitation. Adding 4 uL glycogen to each ethanol precipitation significantly improved the ability to visualize such a small pellet. After ethanol precipitation, the quality of RNA was assessed by tape station.

Isolated RNA was converted to cDNA using the ImProm-II reverse transcription system (Promega) with the RT primer (see table). We also performed reverse transcription on the random RNA library not subjected to U2AF or IP (input sample). We performed PCR on each sample using NEBNext Multiplex Oligos for Illumina (NEB). We assessed PCR product by agarose gel and chose the number of amplification cycles to avoid side products (0 nM-15 cycles, 20 to 320 nM-11 cycles, input-9 cycles). Finally, DNA bands corresponding to about 160 bp were purified from 8% Native PAGE and incubated in 400 uL gel extraction buffer (10 mM Tris, pH 7.0, 300 mM NaCl, 2 mM EDTA) for 1 hour at 65C. DNA was further cleaned using a DNA Clean & Concentrator Kit (Zymo). The correct DNA length (160 bp) was confirmed by tape station.

DNA was pooled into a single tube (67 fMoles per reaction) and sent for deep sequencing at the NCI sequencing facility. Sequencing was performed on a NovaSeq (Illumina) in an S2 flow cell with 2x100 read length. We sequenced to a depth of between 149 million to 450 million reads per binding reaction which allowed us to determine enrichment scores for very long U2AF binding sites. For all samples, at least 90% of reads had a quality score above Q30.

Analysis, performed using the RBNS pipeline (https://github.com/cburgelab/RBNS_pipeline), was too computationally intensive for a single computer. Therefore, the pipeline was performed on Biowulf, a linux cluster maintained by the NIH High Performance Computing center.

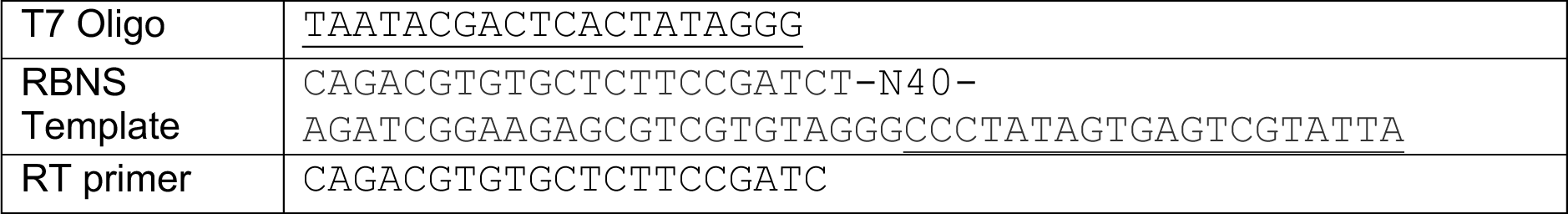

### Predicting relative binding affinities of *in vivo* U2AF binding sites

We randomly selected 2.5 million reads from each U2AF concentration of the RNA Bind-N-Seq assay for analysis in the probound server (http://probound.bussemakerlab.org/) (*20*). Each run also included 2.5 million reads from the input fraction. We generated U2AF binding models for 6mer, 12mer, and 18mer sequences (*see supplementary)*.

We used the 12mer binding model from U2AF = 20 nM to assign affinities to each peak from a previous PAR-CLIP experiment (*6*). We determined the U2AF binding affinity of every 12mer in a single PAR-CLIP peak and assigned the highest affinity value as affinity of that binding site. We did not consider the small fraction of PAR-CLIP peaks that spanned less than 12 nt.

### Competitive Binding Assays

Testing U2AF binding directly to sites identified from PARCLIP would require copious amounts of U2AF. Therefore, we performed competitive binding assays where 50 nM U2AF was prebound to 10 nM Cy5-(U)_12_ RNA and competed off with increasing concentrations of unlabeled competitor RNA. For each titration we measure the IC_50_, the concentration of unlabeled RNA required to compete off 50% of the Cy5-RNA. The relative affinity of two binding sites is the ratio of their IC_50_ values. Binding assays were resolved by EMSA (see EMSA section for details) using the RNA oligos in the table below. While probound predicts a very broad range of affinities, we were only able to validate U2AF binding to sequences with binding affinity greater than 370-fold weaker than the strongest motif. Binding reactions were performed over multiple days. To account for experimental variability, each competitive binding assay was performed in parallel to the ‘3B’ sequence. Reported relative binding affinity is IC_50_ / IC_50 3B Seq_.

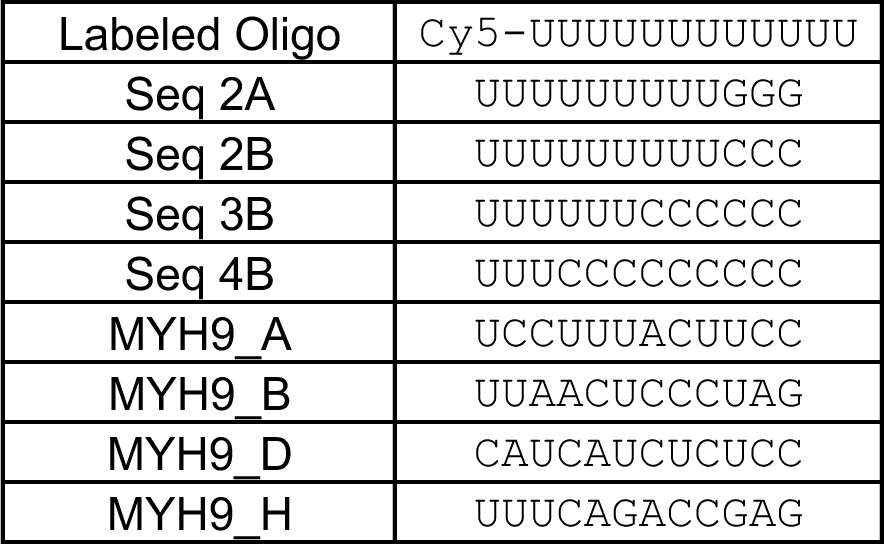

### Ensemble PIFE assay

U2AF binding to 5 nM Cy3-RNA was determined by protein induced fluorescence enhancement (PIFE), in which Cy3 fluorescence increases upon protein binding as previously described (*21*, *40*). Each binding reaction was 60 uL. Cy3 was excited at 510 nm. We measured Cy3 emission measured in 2 nm steps from 550-630 nm. Fluorescence enhancement was greater when the Cy3 was placed after the U2AF1 binding site as opposed to before the U2AF2 binding site. RNA was incubated for at least 5 min with varying concentrations of U2AF in ‘U2AF binding buffer without tRNA’ (25 mM Tris pH 7.0, 150 mM NaCl, 0.5 mM DTT, 10% glycerol, 0.05% Tween 20). Fluorescence spectra were obtained using a Horiba fluorometer and analyzed in MATLAB to (1) remove background autofluorescence from buffer and protein and (2) to determine the relative change in Cy3 fluorescence compared to 0 nM. Background autofluorescence was determined by performing the same fluorescence scan but without RNA (protein + buffer).

### Slide Preparation for smPIFE

A quartz microscope slide (Alfa Aesar) was patterned with 16 small holes to form 8 flow cells (each flow cell has an inlet and and outlet). #1.5 coverslips (Corning) and microscope slides were soaked in toluene, ethanol, and then plasma cleaned. Slides were then incubated in 100 uM mPEG-Si and biotin-PEG-Si (Laysan Bio) overnight in anhydrous toluene. Functionalized quartz slides and coverslips were assembled into microscope flow cells where channels are defined with parafilm. Before each experiment, the flow cell was treated sequentially with 1 mg/ml BSA, 40 ug/ml neutravidin, and biotin-labeled RNA in ‘U2AF binding buffer -tRNA’ + 0.5% Tween-20.

### smPIFE data acquisition

To attach U2AF binding sites to the biotinylated surface, we annealed either the ‘AdML’ or ‘Mutant’ RNA to a DNA handle that was biotinylated on it’s 3’ end. All oligos were purchased through IDT.

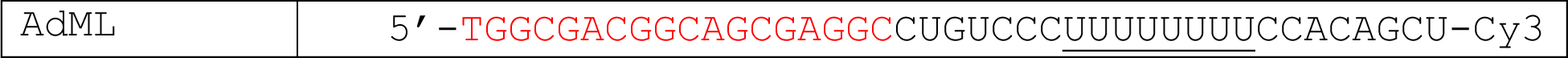
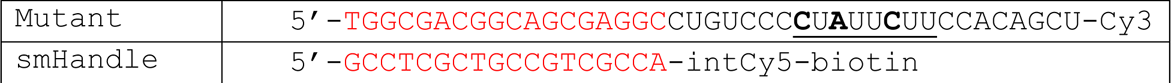

Annealed RNAs were added to the flow cell. After a 5 minute incubation, unbound molecules were removed by a wash with ‘U2AF binding buffer -tRNA’. We then added imaging buffer containing the desired U2AF concentration into the flow cell. Imaging was performed for 500 frames at 1 Hz. The imaging buffer contained 10 mM Tris-HCl (pH 7), 150 mM NaCl, 10% glycerol, 0.5% v/v Tween-20, 0.1 mg/ml BSA, 2 mM Trolox, 0.0115% v/v COT (Cyclooctatetraen, Sigma 138924), 0.012% v/v NBA (3-Nitrobenzyl alcohol, Sigma 146056), 450 ug/ml glucose oxidase (Sigma G2133) and 22 ug/ml catalase (Sigma C3155).

### smPIFE data analysis

In each image stack, the imageJ ‘‘find maxima’’ function was applied to determine the coordinates of each RNA molecule. Time traces were generated by measuring the average intensity of a 3x3 pixel area centered on each molecule throughout the entire stack. We typically identify about 1000 RNA molecules per field of view and attempted to analyze as many RNAs as possible in each video. We ended up analyzing 22 ± 7% and 28 ± 6% of all identified molecules for AdML and Mutant RNAs, respectively. While we attempted to include as many RNAs as possible in the analysis, we discarded traces with multiple photobleaching steps (RNA aggregates), traces that photobleached too quickly (<50 seconds), and uninterpretable traces. Below is a table of the number of molecules identified in each measurement and the number of molecules sent for further analysis:

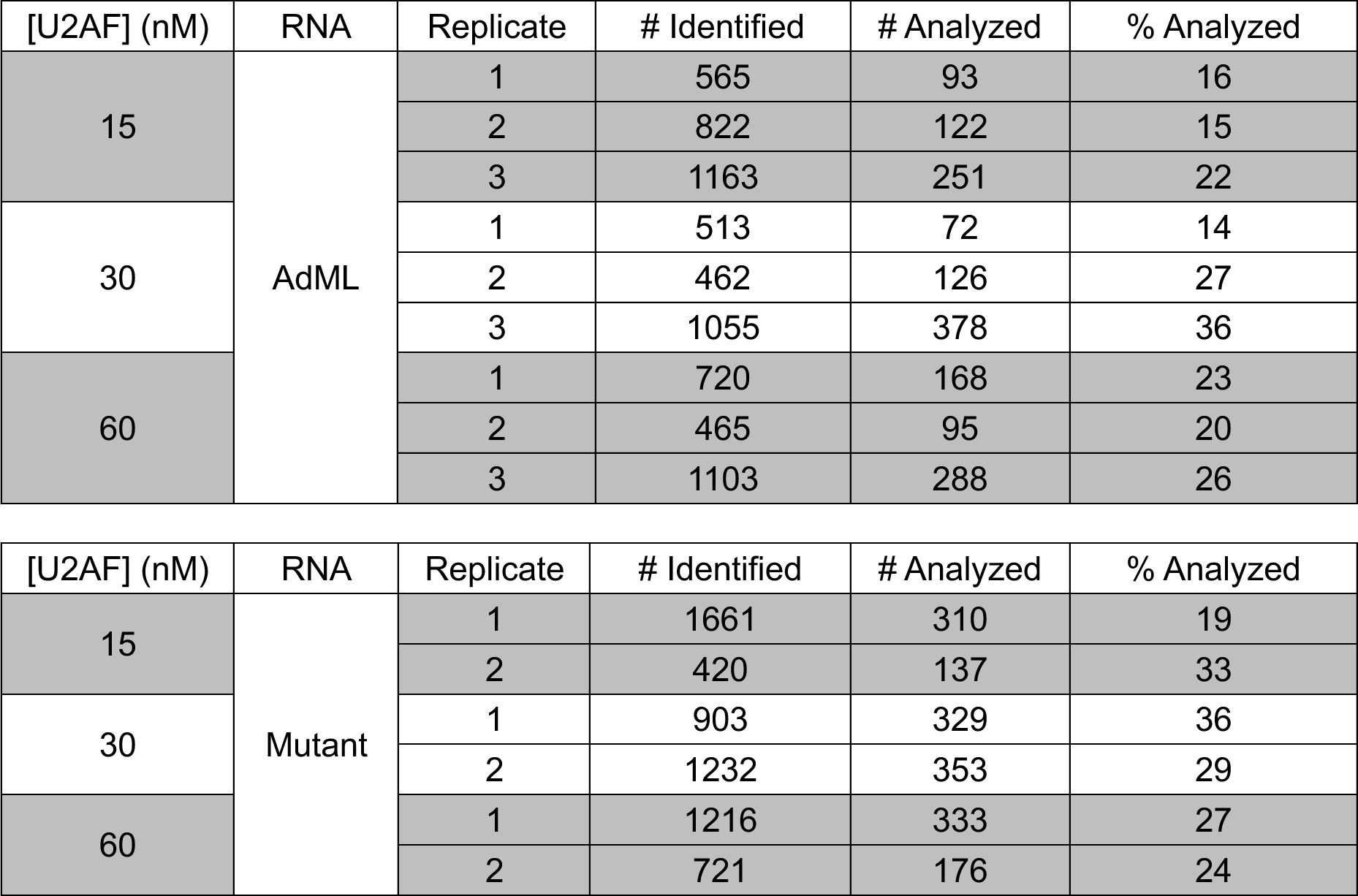

Selected traces were cropped and analyzed using ebFRET (*41*) which infers kinetic parameters based on global HMM fitting. We fit the data to a two state HMM model where the high state represents U2AF in complex with RNA and the low state represents unbound RNA. It becomes difficult to observe binding fluctuations at high concentrations of U2AF where the fluctuations begin to occur very quickly. Nevertheless, we observe a concentration dependent on-rate which indicates the observed fluctuations represent U2AF binding events.

### RNA sequencing

RNA extraction was performed on ∼1 million Human Bronchial Eplithelial Cells (HBEC) using the Qiagen RNeasy mini kit (74106). Genomic DNA and degraded using the Turbo DNase kit (Invitrogen, AM1907). We verified complete depletion of genomic DNA by qPCR. All samples had an RNA integrity value (RIN) above 9.9. We sent 600 ng of each sample for sequencing library preparation by the CCR genomics core which performed rRNA depletion using NEBNext rRNA Depletion Kit v2 (E7400), and sequencing library preparation using NEBNext Ultra II Directional RNA Library Prep Kit for Illumina (E7760). Sequencing was performed using Illumina NextSeq 2000 P3 (200 cycle) reagents at a depth of 163 to 193 million reads per replicate. Raw sequencing files were aligned using Bowtie2 (*42*) which also removed reads corresponding to rRNA and ERCC spike-in sequences (ThermoFisher 4456740). After transcript quantification with Salmon (*43*), we evaluated differential splicing with SUPPA2 (*32*).

### Preparing Cell Lines

We tagged endogenous U2AF2, U2AF1, and H2B with HALO-tags for single-molecule tracking experiments in Human Bronchial Epithelial Cells (HBECs). We used Keratinocyte SFM (Gibco 17005042) for all cell culture. For the U2AF2 cell line, we inserted a HALO-tag on the C-terminus of U2AF2 followed by P2A and a blasticidin marker for selection. For the U2AF1 cell line, we inserted a HALO-tag on the C-terminus of U2AF1 followed by 3XFLAG, T2A, and a blasticidin marker for selection. For orbital tracking assays, we also HALO-tagged the C-terminus of U2AF1 in the MYH9-MS2 cell line created earlier(*6*). Finally, for the H2B cell line, we inserted a HALO- tag on the C-terminus immediately after a TEV site.

Tagging was accomplished with CRISPR-CAS9. Guide RNAs were cloned into the PX458 plasmid to target the C-terminus. Subsequent HALO-tag integration was accomplished by homology directed repair. These are the genomic coordinates of the homology arms:

| Name | Assembly | chr | Left homology arm |  | Right homology arm |  |
| --- | --- | --- | --- | --- | --- | --- |
|  |  |  | Start | End | Start | End |
| U2AF2-HALO | GRCh38.p14 | 19 | 55,673,561 | 55,674,065 | 55,674,069 | 55,674,219 |
| U2AF1-HALO | GRCh38.p14 | 21 | 43,093,105 | 43,093,960 | 43,092,220 | 43,093,098 |
| H2BC8-HALO | GRCh38.p14 | 6 | 26,216,266 | 26,217,066 | 26,215,502 | 26,216,265 |

### Single-molecule tracking

Single-molecule tracking of HALO-tagged proteins was performed with Highly Inclined Laminated Optical sheet (HILO) microscopy on a custom-built microscope. The microscope is built on a Rapid Automated Modular Microscope system (RAMM, Applied Scientific Instrumentation) with synchronized lasers and cameras using an Arduino board. Images are acquired using micromanager (Open imaging). The microscope contains a 100x 1.49 NA oil immersion TIRF objective which corresponds to a pixel size of approximately 150 nm. The automated stage (Applied Scientific Instrumentation) is fitted with an incubator to maintain CO2 and temperature (Oko lab and Tokai Hit). The emission path contains a quad-band dichroic (ZT405/488/561/647 Chroma technology) which is further separated by two long pass filters (T588lpxr, T660lpxr) and emission filters (525/50, 609/58, 736/128, Semrock) and directed onto 3 separate EMCCD cameras (Photometrix).

Tracking was performed in Human Bronchial Epithelial Cells (HBEC) labeled with between 0.5 pM – 1 pM JF646 dye (Promega). At this concentration of dye, each frame contained about 5-10 labeled U2AF or H2B molecules. JF646 was stored at 500 nM in DMSO at -20 C and serially diluted to 0.5 pM in Keratinocyte SFM (Gibco 17005042) (KSFM). After 15 minute incubation at 37C, we removed free dye by washing 3x in KSFM, followed by a 10 minute incubation at 37C, followed by another 3x KSFM wash before imaging.

These experiments are analyzed using the MatlabTrack pipeline (https://github.com/davidalejogarcia/PL_HagerLab.git) (*44*). Slow tracking experiments were conducted at a rate of 1 frame every 3 seconds. During each frame, cells were illuminated for 100 ms with a 647 nm laser (Coherent Obis 1196627) set to power 1 mW (with a 10% neutral density filter). Fast tracking measurements were performed at a rate of 1 frame every 10 ms with constant illumination at laser power set to 15 mW. Tracking was performed in MatlabTrack with the following tracking parameters: filteringèlocal background, no filtering (use SLIMfast), maximum jump è 4, shortest track è 2, gaps to close è 2. The diffusive parameters of a ‘bound’ molecule were determined by tracking H2B-HALO (further analysis è merge and analyze jump histogram) using the approach defined in this paper (*44*). Additionally, because the vast majority of H2B is stably integrated into chromatin, disappearance of an H2B-HALO ‘spot’ corresponds to photobleaching. Therefore, the U2AF dwell time distribution from slow tracking assays was corrected for photobleaching using the approach outlined in this this paper (*44*).

We analyzed fast tracking data in vbSPT (*45*) where tracks were fit to two states, ‘bound’ and ‘unbound’ using the default priors and the following settings:

- time step =0.01
- dim = 2
- trjLin = 2
- bootstrapNum = 10
- fullBootstrap = 0
- init_D = [0.001, 16]
- init_tD = [2, 20]*timestep.

We observe three types of tracks: U2AF is unbound for the entire trajectory, U2AF is bound for the entire trajectory, and U2AF transitions between bound and unbound state during the trajectory. In this very fast imaging condition with high laser power, we suspected that many long trajectories ended because of photobleaching and not becuase diffusion out of illumination volume. Therefore, when determining dwell times at these short time scales, we only analyzed the minority of tracks that show a bound event separated by two unbound events. In this way, we can be sure that we are observing dissociation and not photobleaching.

### Single-molecule FISH

Nascent RNA was visualized by single-molecule FISH (smFISH) as previously described (*6*, *46*). Human Bronchial Epithelial Cells were plated onto #1.5 coverslips, washed 3X with HBSS and fixed by incubating for 10 minutes in 4% EM Grade Paraformaldehyde (Electron Microscopy Sciences) in PBS. After 1X PBS wash, cells were permeabilized by incubating in 70% ethanol at 4C for 1-2 days. Before adding FISH probes, coverslips were incubated 10 minutes in PBS twice and then 5 minutes in wash buffer (10% formamide, 2X SSC). FISH probes were diluted to 50 nM in hybridization buffer (2× SSC (v/v), 10% dextran sulfate (w/v) and 10% deionized formamide (v/v) and added dropwise to parafilm. Coverslips were placed cell-side down in hybridization buffer and incubated at 37C for at least 4 hrs. After incubation, coverslips were incubated at 37C twice for 30 minutes in wash buffer. Coverslips were then washed with 2X SSC followed by PBS. After air drying, coverslips were placed cell side down into mounting media spotted onto microscope slides (Prolong Gold with DAPI, Thermo) and stored at -20C. We used the same probe sets as in Wan et al 2021.

Single-molecule FISH experiments are performed on a custom microscope setup consisting of a RAMM (ASI) chassis and a Zeiss 40x, 1.4 N.A. objective. The emission path is filtered using both a quad bandpass filter and an automated emission filter wheel for imaging DAPI, GFP, Cy3 and Cy5 (VCGR-SPX-P01-PC, Chroma). We’ve installed an automatic stage (ASI) for imaging multiple fields of view using an ORCA-Flash4 V2 CMOS camera (Hamamatsu). The microscope is controlled using micromanager (Open imaging). Image analysis is performed in python (https://github.com/Lenstralab/smFISH_analysis.git).

### 3D Orbital Tracking

Orbital tracking was performed on an inverted Zeiss Observer Z.1 microscope. The microscope contains a Zeiss 63X 1.2 NA water immersion objective and an automated stage (ISS) that is temperature and CO_2_ controlled (Oko Lab and Tokai Hit). U2AF1-HALO (labeled with JF646 (Promega)) was detected with a long pass filter (T647lpxr) followed by a bandpass filter (ET706/95m) (Chroma). The GFP channel is filtered using a ET525/50m filter (Chroma). Data is collected with avalanche photodiodes (Excelitas). Tracking was performed using SIM FCS software (ISS).

We selected cells with visible MS2 spots for imaging. These represented cells actively transcribing MYH9 pre-mRNA. Based on single-molecule FISH measurements, the majority of MYH9 transcription sites produce 1-2 pre-mRNAs per burst (**figure S2H**). Therefore, when tracking begins, we typically observe only 1-2 downward steps in MYH9 fluorescence (**Figure S2K**) which corresponds to splicing events. Furthermore, very few traces show increases in fluorescence which corresponds to transcription. Since we begin tracking only after transcription of the stem-loops, we begin the measurement with no understanding of where we are in the lifetime of the intron. For this reason, we are unable to measure the total time to splice by autocorrelation as previously (*27*, *28*). However, cross-correlation, which relates the timing of U2AF binding during the lifetime of the pre-mRNA, shows a peak near 0 minutes which means MYH9 splicing and U2AF binding overlap. Furthermore, perturbation (whether DDX42 knockdown or pladienolide B treatment) broadens the cross-correlation curve indicating slowed splicing relative to the unperturbed control.

Cells were labeled with 5-10 pM JF646 for 15 minutes at 37C. While this is 10 to 20-fold higher dye concentration than the SMT measurements, many U2AF molecules will still go unlabeled. However, increasing the dye concentration further adds to the background fluorescence signal. Therefore, we determined that this dye concentration was best suited to observe single U2AF bindning events while minimizing background signal. Not all traces show clear binding fluctuations in the U2AF channel (**figure S2K**), likely due to underlabeling.

While these measurements indicate that U2AF association to MYH9 occurs about 8 minutes after transcription of the stem loops, it is important to note that because this assay requires underlabeling of U2AF the actual initial time for U2AF to bind is likely much faster. However, we reason that since we use the same dye concentration for all assays, this result reflects the relative change in spliceosome assembly after perturbation.

### Mass-spectrometry

Human Bronchial Epithelial Cells (HBECs) expressing endogenously tagged U2AF1-3XFLAG underwent subcellular fractionation as previously described (*47*). In addition, a parental HBEC line lacking the endogenous U2AF1 tag was prepared to serve as a control. This control was critical for identifying nonspecific interactions, ensuring that observed binding events were attributable to the specific interactions of U2AF1-3XFLAG. The experiment was conducted with three biological replicates per sample group, each consisting of approximately 20 million cells. Briefly, cells washed with PBS at room temperature were detached using trypsin, collected in ice-cold PBS, and transferred to pre-chilled 15 ml conical tubes to inhibit degradation. After centrifugation at 500 x g for 5 minutes at 4°C, the supernatant was removed, and the cell pellet weighed. The pellet was then washed with ice-cold 1X PBS and resuspended in hypotonic lysis buffer (HLB) at a ratio of 1 ml per 75 mg of cells, with a 1X final concentration of protease inhibitor (PI) cocktail and phosphatase inhibitor (PhI) solution. Following a 10-minute incubation on ice with intermittent vortexing, the suspension was centrifuged at 800 x g for 10 minutes to collect the cytoplasmic fraction. The nuclear pellet underwent four washes with HLB before being resuspended in nuclear lysis buffer (NLB) at half the initial HLB volume, with PI and PhI added. Nuclei were then sonicated ten times, alternating between 30 seconds of sonication and 30 seconds of cooling. Subsequent centrifugation at 18,000 x g for 15 minutes at 4°C clarified the nuclear extracts.

For immunoprecipitation (IP), 30 µL of magnetic beads (specific for FLAG-tag affinity) were prepared following the manufacturer’s instructions (Sigma M8823), washed with NLB, and incubated with 500 µL of nuclear extract overnight at 4°C with rotation to ensure thorough binding. Afterward, beads were washed three times with NLB to remove nonspecifically bound components.

### On-bead digestion and TMTpro labeling

Samples were digested on bead by treatment with 50µL each of lysis buffer, reducing buffer, and alkylation buffer provided with the EasyPep kit (Thermo #A40006) then incubated at 25°C 1.5hrs in the dark. Samples were treated with 20µL of 100ng/µL trypsin/LysC and incubated at 37°C overnight for 20hrs with shaking at 1000rpm. Supernatant was then moved to a clean tube and treated with 50µL of 5µg/µL TMTpro in 100%ACN and incubated at 25°C for 2hrs. Samples were then quenched with 50µL of 5% hydroxylamine, 20%FA and incubated for 20 minutes before combining samples within each plex. Each plex was cleaned using EasyPep mini spin columns (Thermo #A40006) and drying eluted peptides by speed-vac.

### LC-MS/MS analysis

Each plex was loaded twice onto a Dionex U3000 RSLC in front of a Orbitrap Eclipse (Thermo) equipped with an EasySpray ion source Solvent A consisted of 0.1%FA in water and Solvent B consisted of 0.1%FA in 80%ACN. Loading pump consisted of Solvent A and was operated at 7 μL/min for the first 6 minutes of the run then dropped to 2 μL/min when the valve was switched to bring the trap column (Acclaim™ PepMap™ 100 C18 HPLC Column, 3μm, 75μm I.D., 2cm, PN 164535) in-line with the analytical column EasySpray C18 HPLC Column, 2μm, 75μm I.D., 25cm, PN ES902). The gradient pump was operated at a flow rate of 300nL/min and each run used a linear LC gradient of 5-7%B for 1min, 7-30%B for 133min, 30-50%B for 35min, 50-95%B for 4min, holding at 95%B for 7min, then re-equilibration of analytical column at 5%B for 17min. All MS injections employed the TopSpeed method with three FAIMS compensation voltages (CVs) and a 1 second cycle time for each CV (3 second cycle time total) that consisted of the following: Spray voltage was 2200V and ion transfer temperature of 300 ⁰C. MS1 scans were acquired in the Orbitrap with resolution of 120,000, AGC of 4e5 ions, and max injection time of 50ms, mass range of 350-1600 m/z; MS2 scans were acquired in the Orbitrap using TurboTMT method with resolution of 15,000, AGC of 1.25e5, max injection time of 22ms, HCD energy of 38%, isolation width of 0.4Da, intensity threshold of 2.5e4 and charges 2-6 for MS2 selection. Advanced Peak Determination, Monoisotopic Precursor selection (MIPS), and EASY-IC for internal calibration were enabled and dynamic exclusion was set to a count of 1 for 15sec. The only difference in the methods was the CVs used, one method used CVs of -45, -60, -75 and the second used CVs of -50, -65, -80.

### Database search and post-processing analysis

Raw files from each plex were batched together as fractions and searched with Proteome Discoverer 2.4 using the Sequest node. Data was searched against the Uniprot Human database from Feb 2020 using a full tryptic digest, 2 max missed cleavages, minimum peptide length of 6 amino acids and maximum peptide length of 40 amino acids, an MS1 mass tolerance of 10 ppm and MS2 mass tolerance of 0.02 Da. Variable modifications of oxidation on methionine (+15.995 Da) and TMTpro (+304.207) on lysine and peptide N-terminus as well as fixed carbamidomethyl on cysteine (+57.021). Percolator was used for FDR analysis and TMTpro reporter ions were quantified using the Reporter Ion Quantifier node and normalized on total peptide intensity of each channel. Proteins were filtered using a FDR cutoff of 1% and have quantitative values in at least 4 samples to be included in the final list. An absolute log2 fold change of 0.6 and pvalue of 0.05 were used to determine if a protein/phosphopeptide was differentially regulated. TMTpro channel assignment were:

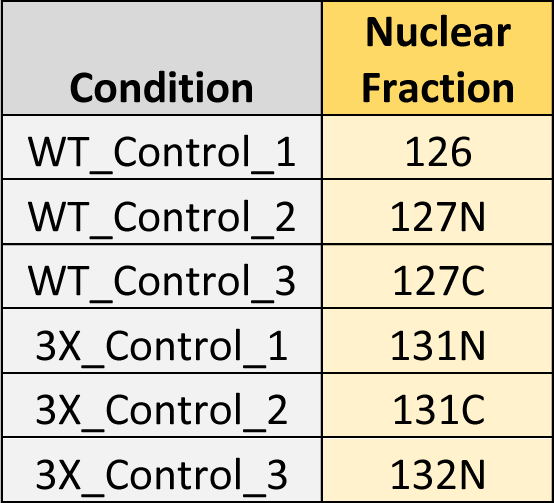

The list of protein in each snRNP were complied using the ‘class_family’ complumn from spliceosome database (*38*) (U2 Associated: “17S U2 snRNP associated,” U2: ‘’17S U2 snRNP’’, U1: “U1 snRNP,” U4/U6 & U5 snRNP: combined “U4/U6 snRNP”, “U5 snRNP” & “tri-snRNP”).

### DDX42 knockdown

DDX42 knockdown was performed by electroporation with the neon kit (Invitrogen). We electroporated 500K cells with 20 pMoles of two separate siRNAs pools (Invitrogen silencer AM16708 and Dharmacon siGENOME smartpool M-012393-00-0005) into 120 uL Buffer R (Invitrogen) for a final concentration of 333 nM siRNA. Cells were electroporated with 4-10 ms pulses at 1230 volts. Optimal siRNA concentration was determined by titration (**figure S6A).** For control assays, we used 333 nM siGENOME non-targeting siRNA pool #1 (D-001206-13-05, Dharmacon). Knockdown efficiency was assessed by western blot using a DDX42 polyclonal antibody from Invitrogen (PA5-54828) 3 days post electroporation. For RNA sequencing experiments, this reaction was scaled up to 750K cells per DDX42 knockdown.

## SUPPLEMENTARY FIGURES

**Fig S1.**
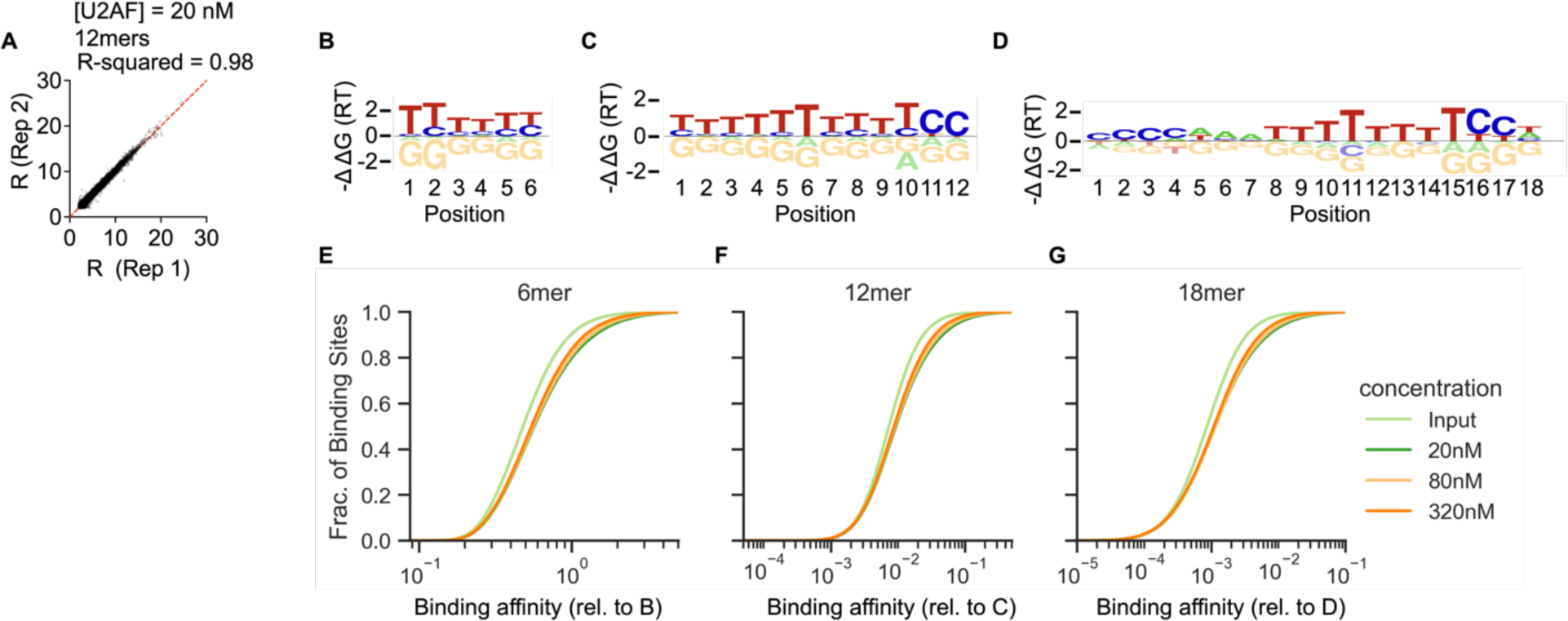
Details on RBNS and Probound Assays **A)** RBNS was performed with two replicates of 0, 20, 80, 320 nM U2AF. Comparing enrichment (R) for significantly enriched kmers (Z=3) for U2AF = 20 nM indicates strong agreement between two measurements. **B)** U2AF binding model from probound for 6mers. **C)** U2AF binding model from probound for 12mers. **D)** U2AF binding model from probound for 18mers. **E)** Cumulative distribution of predicted binding affinity from the 6mer Probound model for enriched RNA in different replicates of the RBNS assay (input, 20 nM, 80 nM, 320 nM). The RNAs enriched for 20 nM U2AF have the highest binding affinity. Reads from the the initial library (input) have the lowest binding affinity. **F)** Same as (E) but for 12mer probound model. **G)** Same as (E) but for 18mer probound model.

**Figure S2.**
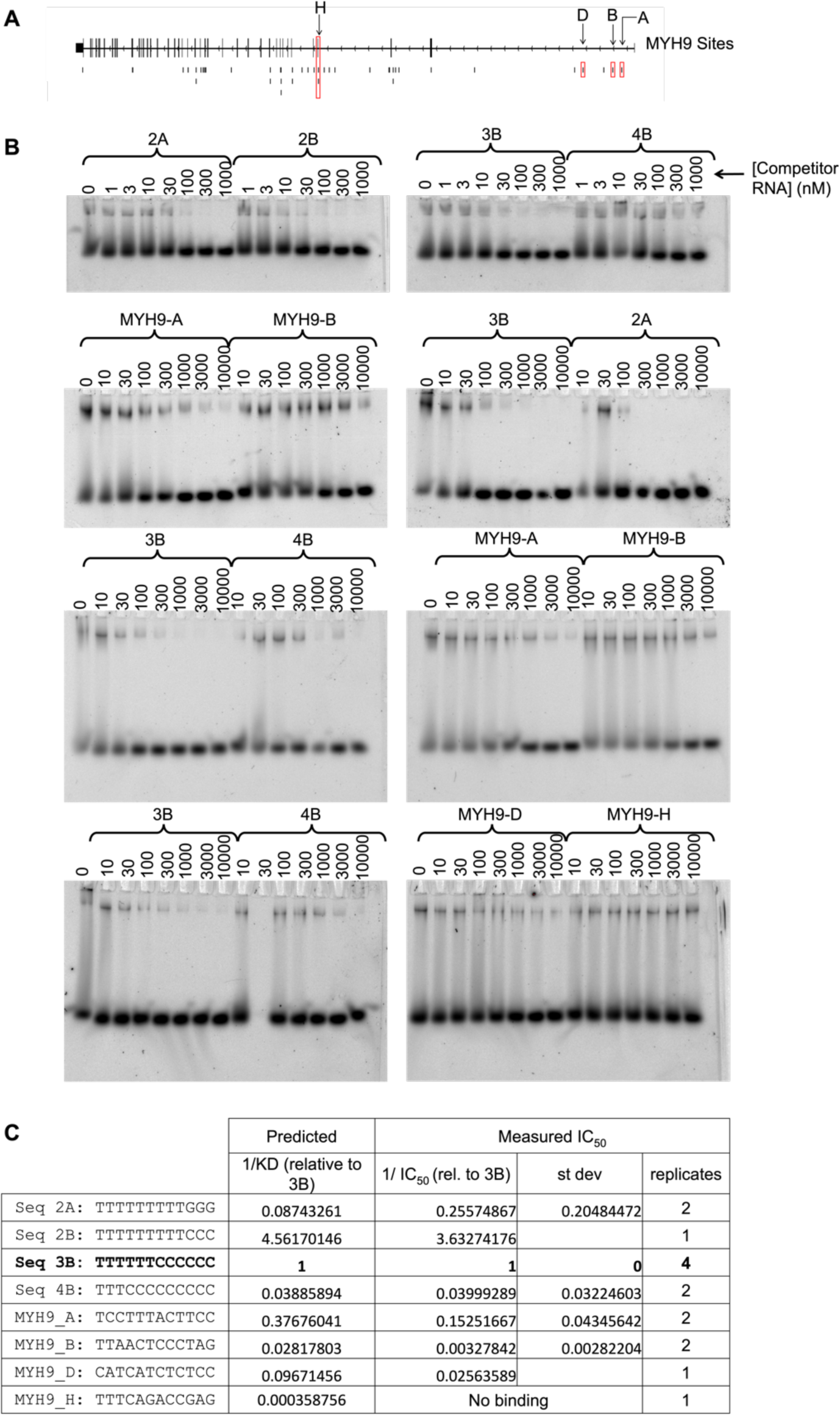
Details on Probound validation by competitive binding assay. **A)** We tested U2AF binding to various binding sites identified in PAR CLIP in the MYH9 pre- mRNA. **B)** EMSA measurements from competitive binding assays. **C)** Comparing predicted relative binding affinity from Probound 12mer model with measured U2AF binding affinity from EMSA measurements.

**Figure S3.**
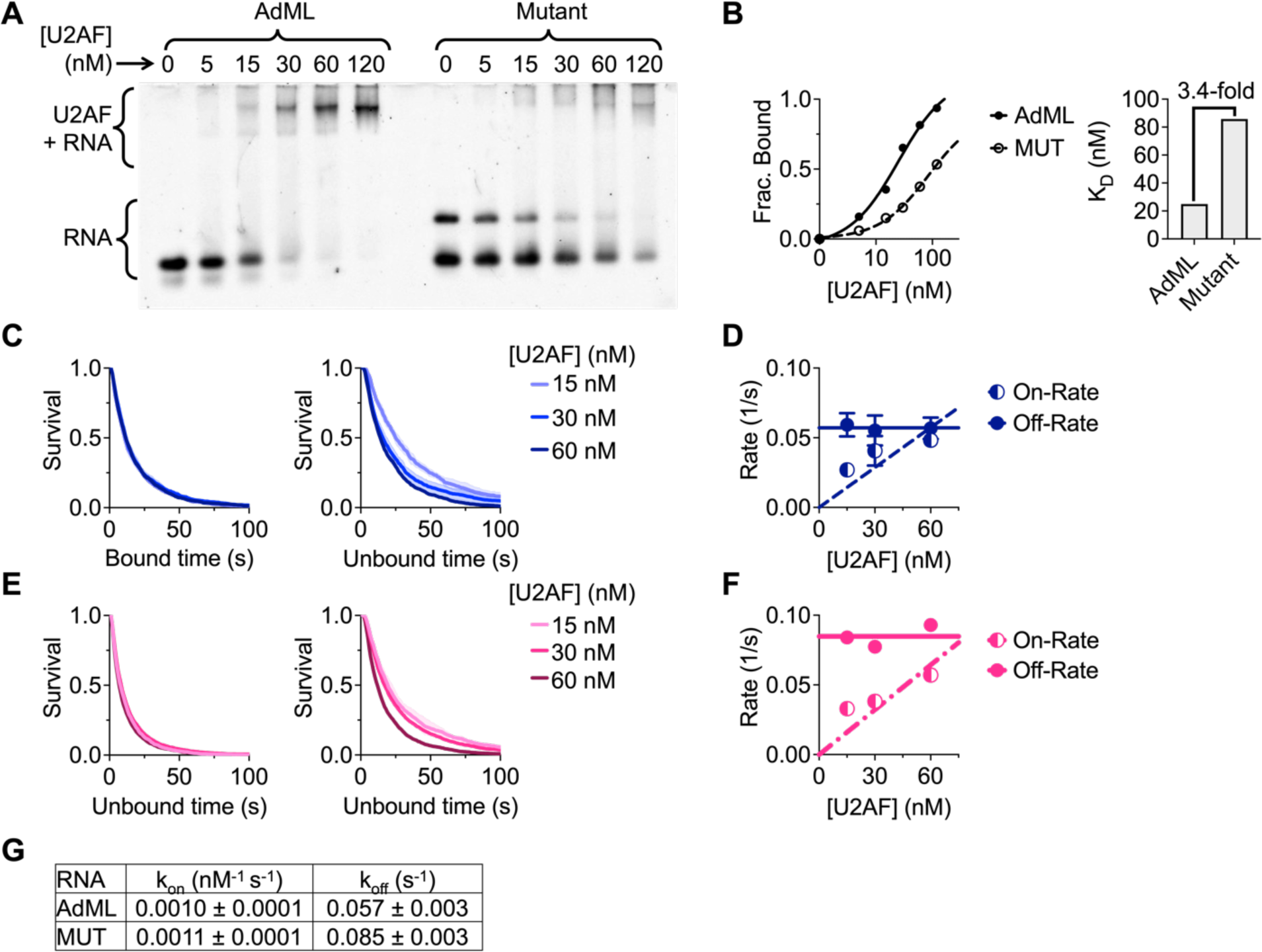
Details on in vitro binding assays. **A)** Measuring U2AF binding affinity to AdML and Mutant 3’SSs by EMSA. **B)** Quantification of EMSA measurement reveals a 3.4-fold difference in binding affinity (n=1 replicate). **C)** Dwell time distributions of U2AF in the bound state and unbound state for 3 concentrations of U2AF binding to the AdML 3’SS (n=3 replicates for each concentration). **D)** Dwell time distributions were fit to a single exponential distribution to determine the off-rate (from bound time survival plot) and on-rate (from unbound time survival plot) for U2AF binding the AdML 3’SS. This plot shows both rates at each concentration of U2AF. The on-rate is concentration dependent and the off-rate is concentration independent. **E)** Same as (C) but for the mutant 3’SS. **F)** Same as (D) but for the mutant 3’SS. **G)** Table of measured rates of U2AF binding to both 3’SSs.

**Figure S4.**
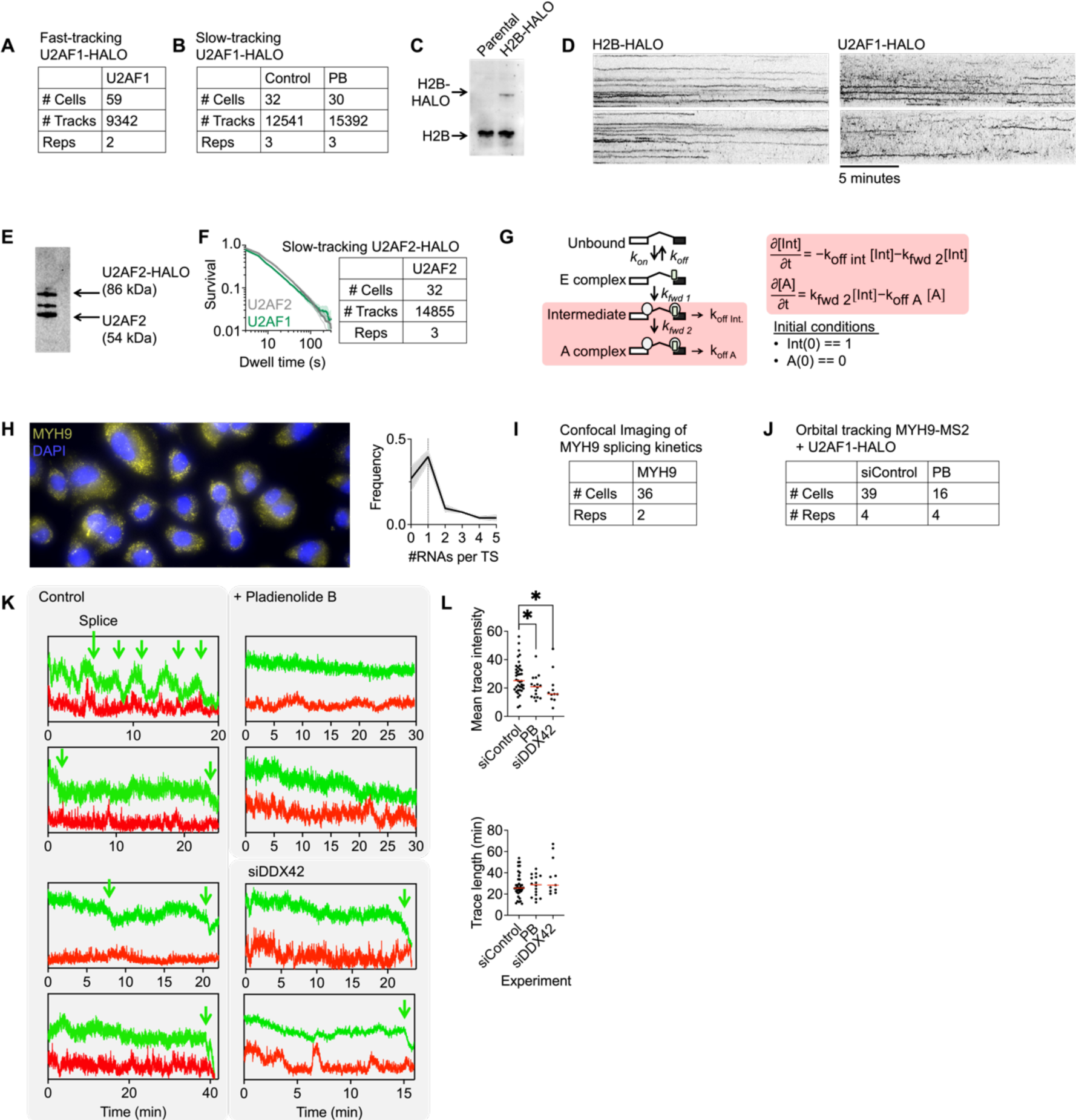
Live cell single-molecule tracking of U2AF in spliceosome assembly. ***A)*** Experimental details for fast tracking dataset (100 Hz acquisition rate, 10 seconds) for U2AF1-HALO. ***B)*** Experimental details for slow tracking dataset (0.33 Hz acquisition rate, 20 minutes) for U2AF1-HALO (with and without PB treatment). ***C)*** Western blot of H2B-HALO HBEC cell line. ***D)*** Kymographs of H2B-HALO and U2AF1-HALO in slow tracking conditions. ***E)*** Western blot of U2AF2-HALO HBEC cell line. ***F)*** Left: Survival plot comparing U2AF1-HALO and U2AF2-HALO dwell times in slow SMT assay. Dwell times appear similar for both cell lines. Right: Experimental details for U2AF2-HALO slow SMT dataset. ***G)*** Binding model used to interpret slow SMT data. Equation 1 comes from master equation with initial conditions. ***H)*** Single-molecule FISH measurements using probes that target the 3’UTR of MYH9. Histogram of number of RNAs per transcription site reveals MYH9 is actively transcribed in many cells. ***I)*** Experimental details for overnight imaging of MYH9 transcription and splicing kinetics. ***J)*** Experimental details for orbital tracking measurements for cells treated with and without pladienolide B (PB). ***K)*** Raw time traces from orbital tracking measurements of MYH9-MS2 (green) and U2AF1- HALO (red). This assays requires U2AF1-HALO to be underlabeled with 5-10 pM JF646 dye. For this reason, while splicing appears in most tracks (green arrows), we did not observe clear U2AF binding in every cell. See methods for details. ***L)*** Plots of mean trace intensity and trace length for each cell analyzed. Generally speaking, transcription sites were followed for more than 20 minutes for all three conditions.

**Fig. S5.**
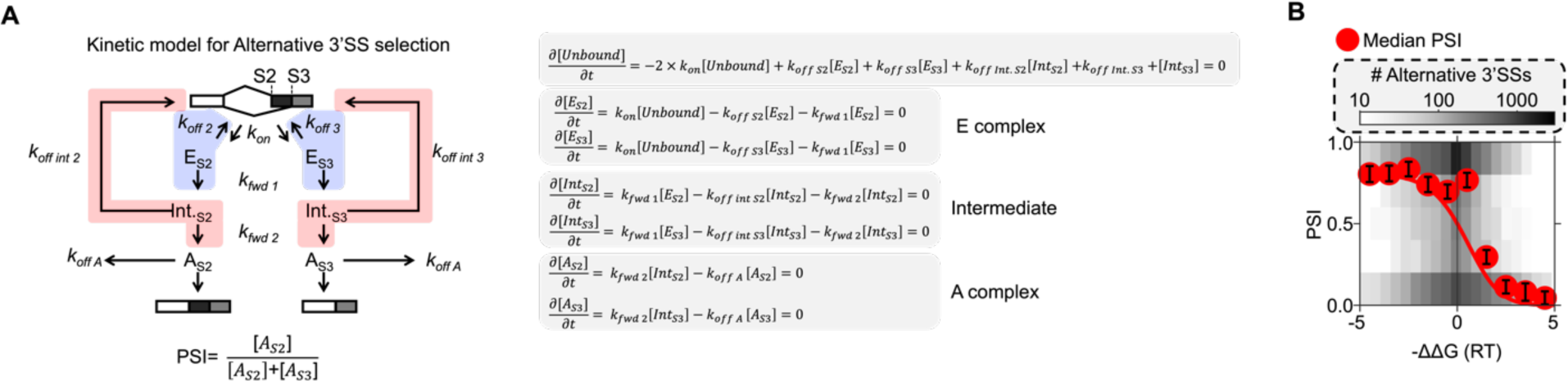
Alternative splicing model A) Schematic and master equations for alternative splicing model (equation 2). B) Plotting median PSI from alternative splicing analysis as in figure 3D. PSI values were binned by −ΔΔG, the free energy of U2AF binding the S3 site relative to the S2 site. Red circles represent median PSI of each bin. Error bars represent the average standard deviation of PSI for each alternative 3’SS (n = 2 replicates). The red curve represents a fit to the alternative splice model (equation 2) (K_s_ = 0.288 ± 0.001, K_f_ = 10 (not fit)). To fully describe the variability of this data, we have also included a binned scatter plot showing the underlying variability. The color of each bin corresponds to the number of 3’SSs with contained with a particular −ΔΔG and PSI combination as indicated by scale bar.

**Figure S6.**
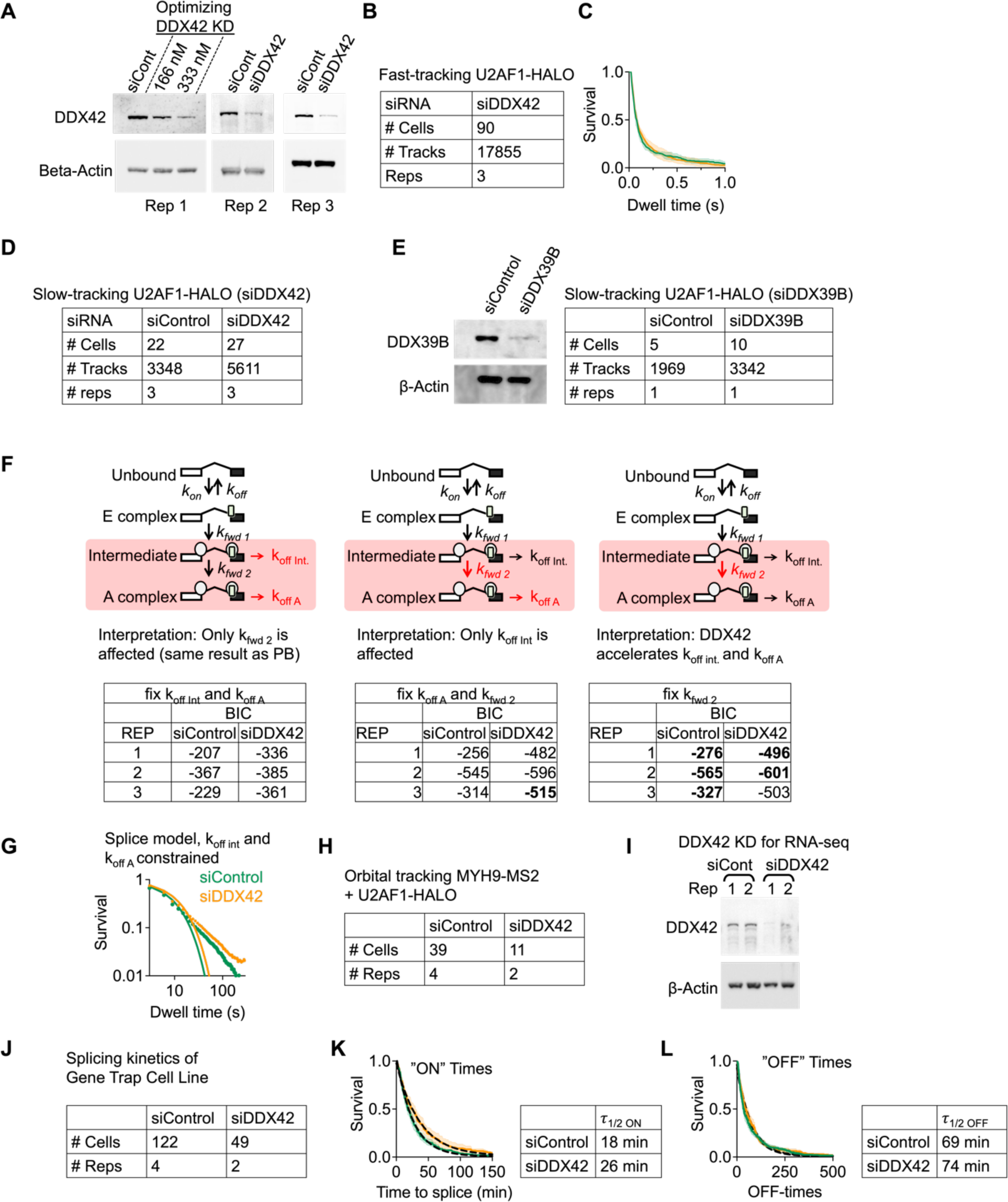
Characterizing the role of DDX42 in spliceosome assembly kinetics **A)** Western blot of DDX42 knockdown for each replicate of the SMT assay **B)** Experimental details for fast tracking of U2AF1-halo after DDX42 knockdown. **C)** Dwell times times measured in fast SMT assay for siControl and siDDX42 experiments. DDX42 does not seem to impact the U2AF1-HALO dwell times on these time scales. **D)** Experimental details for slow tracking of U2AF1-HALO after DDX42 knockdown. **E)** Western blot of DDX39B knockodown and experimental details on slow tracking of U2AF1-HALO after DDX39B knockodown. **F)** Model testing for simultaneous fitting of siControl and siDDX42 slow SMT datasets. We first tried the exact same fitting conditions as the PB experiment (left panel). Fitting was improved by constraining k_fwd 2_ and k_off A_ (middle panel) and further improved by only constraining k_fwd 2_ (right panel). Results of model testing are shown in table for each replicate. Best fits (as indicated by Bayesian Information Criterion values) are indicated in bold. **G)** Simultaneous fit for siControl and siDDX42 slow SMT datasets to the same model as used for PB experiment. We deemed this fit inadequate. BIC values are shown in the left panel of F. **H)** Experimental details for orbital tracking experiment (siControl and siDDX42). **I)** Western blot of DDX42 knockdown for sample sent for RNA-sequencing. **J)** Experimental details for overnight confocal imaging of gene trap cell line. **K)** Comparison of ON-time distributions for siControl and siDDX42 in the polyclonal gene trap cell line. ON-time represents the splicing kinetics. **L)** Comparison of OFF-time distributions for siControl and siDDX42 from imaging the polyclonal gene trap cell line. ‘OFF-time’ distribution represents the time between a splicing event and the start of a transcription event.

